# The regulatory grammar of human promoters uncovered by MPRA-trained deep learning

**DOI:** 10.1101/2024.07.09.602649

**Authors:** Lucía Barbadilla-Martínez, Noud Klaassen, Vinícius H. Franceschini-Santos, Jérémie Breda, Miguel Hernandez-Quiles, Tijs van Lieshout, Carlos G. Urzua Traslaviña, Hatice Yücel, Minh Chau Luong Boi, Celia Hermana-Garcia-Agullo, Sebastian Gregoricchio, Wilbert Zwart, Emile Voest, Lude Franke, Michiel Vermeulen, Jeroen de Ridder, Bas van Steensel

## Abstract

One of the major challenges in genomics is to build computational models that accurately predict genome-wide gene expression from the sequences of regulatory elements. At the heart of gene regulation are promoters, yet their regulatory logic is still incompletely understood. Here, we report PARM, a cell-type specific deep learning model trained on specially designed massively parallel reporter assays that query human promoter sequences. PARM requires ∼1,000 times less computational power than state-of-the-art technology, and reliably predicts autonomous promoter activity throughout the genome from DNA sequence alone, in multiple cell types. PARM can even design purely synthetic strong promoters. We leveraged PARM to systematically identify binding sites of transcription factors (TFs) that are likely to contribute to the activity of each natural human promoter. We uncovered and experimentally confirmed striking positional preferences of TFs that differ between activating and repressive regulatory functions, as well as a complex grammar of motif-motif interactions. For example, many, but not all, TFs act as repressors when their binding motif is located near or just downstream of the transcription start site. Our approach lays the foundation towards a deep understanding of the regulation of human promoters by TFs.

**Highlights:**

- Causality-trained deep learning model PARM captures regulatory grammar of human promoters
- PARM is highly economical, both experimentally and computationally
- Transcription factors have different preferred positions for their regulatory activity
- Many (but not all) transcription factors act as repressors when binding downstream of transcription start sites

## INTRODUCTION

### Human promoters explain nearly half of genome-wide expression

Promoters are the core regulatory elements of all genes. Their activity ensures the correct transcription level of each individual gene, which is critical for homeostasis and response to a wide diversity of signals. To a large extent, promoter activity is determined by a multitude of transcription factors (TFs) that bind to short DNA sequence motifs near the transcription start site (TSS). A key challenge is to construct computational models that can predict promoter activity from DNA sequence. It was previously estimated that the autonomous activity of promoters (∼1 kb upstream of the TSS) explains ∼50% of the genome-wide variance in transcription in human cells^1,2^. From this one may infer that the remaining ∼50% of variance can be attributed to the combined effects of other features such as distal enhancers, insulator elements, DNA methylation, chromatin packaging, and random technical or biological noise. This implies that promoter sequences play a larger role than any other individual mechanism in determining global gene activity patterns. Moreover, promoters harbour by far the most regulatory potential per kb of genomic sequence because they only cover ∼1% of the human genome, while an estimated ∼1 million putative enhancers and other regulatory elements collectively cover ∼10-30% of the genome^3^. Furthermore, human non-coding variants linked to diseases and complex traits are particularly enriched in promoters^4,5^. We therefore focus here on the regulatory logic of promoter sequences.

### Predictions by deep learning

Deep-learning (DL) models that predict human gene activity from DNA sequences have so far utilised large sets of epigenome mapping data as training input^6–9^. This has relied on the assumption that epigenome features such as specific histone modifications often mark active regulatory elements^3^. However, epigenome maps tend to be imperfect predictors of regulatory element activity^10^ and can be confounded by long-range auto-correlative patterns^11^. More recently, DL models trained on transcriptome data have been used to identify DNA features that drive transcription initiation^2,12,13^. Importantly, both epigenome maps as well as transcriptomics data are inherently correlative and do not identify *causal* links between promoter sequence and activity. Moreover, in both cases training data is limited to human DNA sequences in their endogenous context, which may represent insufficient sequence variation to inform complex DL models^14^.

### MPRAs can provide causality information

Massively parallel reporter assays (MPRAs) offer an alternative source of training data. Here, millions of genomic DNA sequences, each several hundred bp long, are tested systematically for their autonomous regulatory activity in a reporter assay. Because each fragment is unique and only partially overlaps with other fragments in the dataset, an enormous increase in meaningful training data variability is achieved. Moreover, because each fragment is tested in isolation, measured activity can be unambiguously assigned to this fragment, and hence inference of the causal roles of specific DNA sequences is expected to be more straightforward than with epigenome maps, particularly when a large number of partially overlapping fragments are probed (**Figure 1a**). After early explorations^15^, evidence that MPRAs in combination with DL can yield powerful predictive models has been reported for *Drosophila* and human enhancers^14,16^ and yeast promoters^17^ but has not yet been used to functionally dissect the grammar of mammalian promoters.

**Figure 1:**
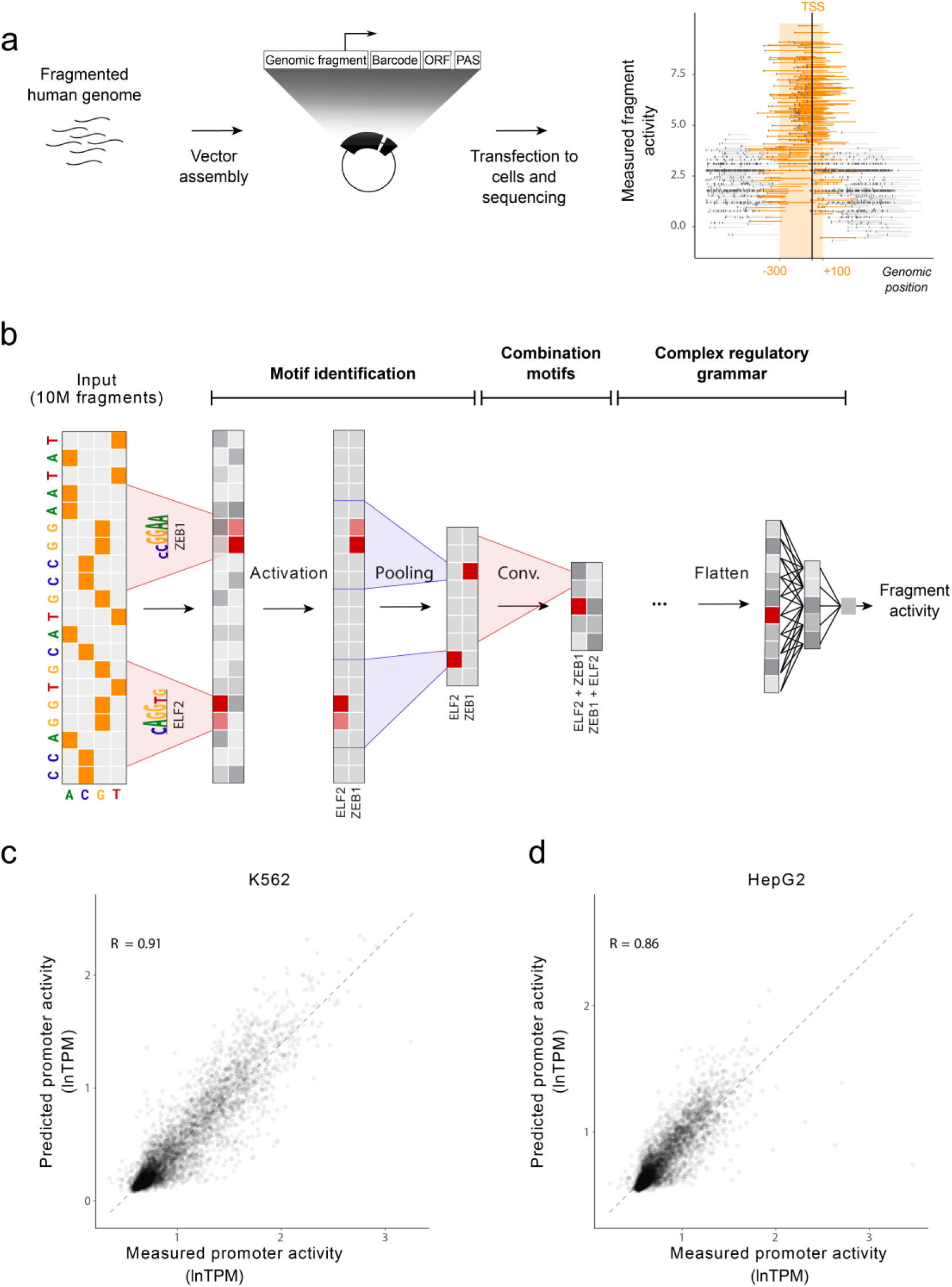
PARM methodology. **a,** Structure of training data. The autonomous promoter activity of millions of random fragments from the human genome were tested by MPRA using a barcoded reporter design^18^. Only fragments that overlap with a window −300 to +100 bp relative to known transcription start sites (TSSs) were selected for training. ORF: Open Reading Frame, PAS: PolyAdenylation Signal. **b,** PARM utilises a CNN that comprises multiple layers. As input, one-hot encoded sequences of all tested genomic fragments are used together with the MPRA-measured promoter activities. In the initial layers, the network learns simpler patterns resembling motifs recognized by Transcription Factors (TFs). Subsequent layers of the network learn more complex regulatory grammar like interactions of TFs. The model output is the predicted promoter activity of any input sequence. **c-d,** Pearson correlation between predicted and measured activity of 5,204 human promoters that were randomly left out from the training data. Promoter activity is according to MPRA measurements in K562 and HepG2 cells^18^, in transcripts per million (TPM), averaged across all overlapping fragments.

### Scope and main findings

Here, we used MPRA-trained DL to build computational models that reliably predict human cell-type specific promoter activity from 600 bp of promoter-proximal DNA sequence alone. These models, which we named Promoter Activity Regulatory Model (PARM), enable in silico study of the regulatory potential and mutational consequence of sequences at a scale not possible *in vitro*, leading to tangible hypotheses that can be validated in the lab. For instance, we leveraged PARM to systematically identify the sequence elements that likely contribute to the transcriptional activity across all human promoters, and uncover the probable TFs that bind to these elements. We also deduce roles for novel sequence elements, and for one example identify the cognate TF. From this rich resource, we deduce cell-type specific and shared mechanisms of TF action across human promoters. PARM infers the activity of TFs as activators and repressors, and often both. Repressive and activating functions of many TFs show striking positional preferences relative to the TSS. Most prominently, many TFs act as repressors if bound downstream of the TSS, suggesting that they may act as roadblocks for RNA polymerase. Furthermore, the activity of some TFs can be strongly dependent on the baseline activity of the promoter, and the model includes detailed combinatorial logic of TFs. PARM provides a foundation to unravel the complex regulatory logic of human promoters at unprecedented scale and resolution.

## RESULTS

### Construction of MPRA-trained DL models of promoter activity

#### Input dataset

We initially set out to construct a sequence-based DL model of promoter activity by training it on data from an MPRA that probes autonomous promoter activity of millions of genomic DNA fragments (**Figure 1a**). These MPRA data were generated in K562 human erythroleukemia and in HepG2 hepatocellular carcinoma cell lines^18^. Crucially, each position in the genome is covered on average by ∼240 random, partially overlapping fragments (**Figure 1a**, right panel), enabling the construction of a DL model as outlined above. For training of the model, we selected a total of 10,403,840 probed fragments of 88-600 bp length that overlapped by at least one nucleotide with a window positioned from −300bp to 100bp around a curated set of 30,607 transcription start sites (TSS) (See Methods).

#### PARM design

PARM consists of a convolutional neural net (CNN), which has previously been used to model similar data structures^15–17,19^. We extensively optimised the precise CNN architecture and hyperparameters (**Figure 1b; Figure S1**; see Methods). No prior knowledge other than the MPRA data was used for training the model. MPRA data from each cell line was used separately to train a cell type specific model.

#### Accurate prediction of autonomous promoter activity

Strikingly, in a cross-validation approach, we found that the trained PARM could predict with high accuracy the measured activity of 5,204 promoters that were randomly omitted from the training and validation set (R = 0.91 for K562 and R = 0.86 for HepG2; **Figure 1c-d**). Note that the experimental replicate measurements of these promoter activities correlate on average with R = 0.99 and 0.93, respectively, imposing upper limits to the estimates of this accuracy in the case of HepG2.

#### PARM can design fully synthetic human promoters

As a stringent test of PARM, we investigated whether it could be used to automatically generate synthetic human promoters, starting from purely random sequences. For this, we adapted a previously reported genetic algorithm^17^. Briefly, the algorithm started with 200 purely random sequences of 232 nucleotides. These sequences were then randomly recombined and peppered with random point mutations; from the resulting sequences, the 200 with the highest promoter activity predicted by PARM were selected to compose a new generation (**Figure 2a**). This cycle of mutagenesis and selection was repeated in silico up to 500 times, which invariably yielded a diversity of sequences that were predicted to be highly active promoters (**Figure 2b**). To verify these predictions, we synthesised 20 of these sequences, cloned them into a barcoded reporter vector and experimentally determined their relative activity in a multiplexed reporter assay. As reference, we tested in parallel 27 equally sized fragments derived from the most active natural human promoters (**Figure 2c**). The synthetic promoters were predicted to be more active than these potent natural promoter sequences (**Figure 2d**). The reporter assays confirmed that virtually all predicted synthetic promoters were active, and on average even more active than the reference set (**Figure 2e**). Sequence analysis of the synthetic promoters indicated that they typically harbour motifs of known activator TFs in K562 cells such as ELK, CREB1 and ATF factors, yet each of these promoters carries a different combination of motifs (**Figure 2f**).

**Figure 2:**
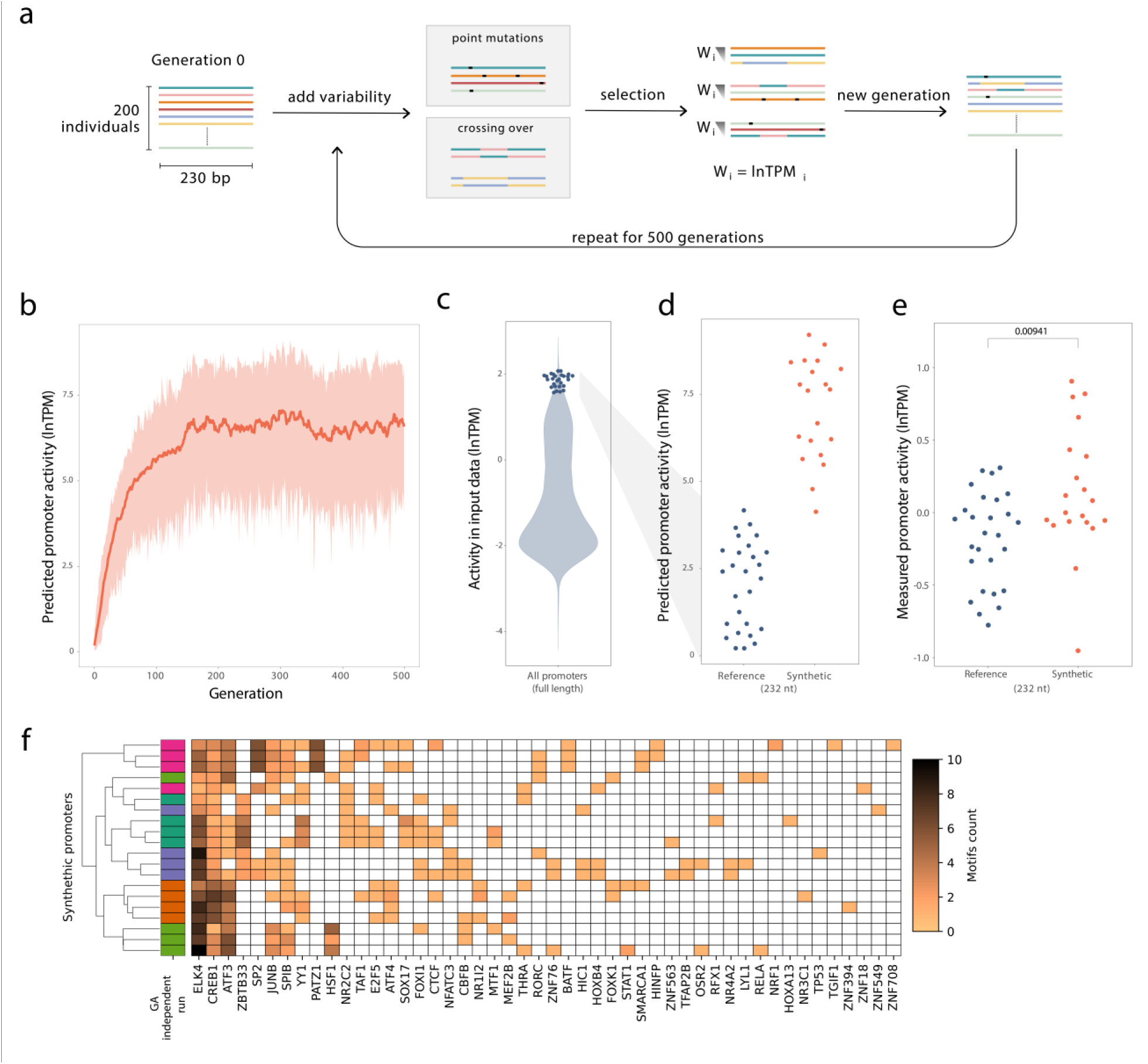
PARM can design highly active synthetic promoters. **a,** Schematic workflow of the genetic algorithm (GA), adapted from^17^. An initial population of 200 random sequences (230 bp each) initiates the GA. The algorithm adds variability to those via point mutations and crossing over (probability of an individual receiving a point mutation Pμ = 0.8; probability of two individuals crossing over Pχ = 0.5). The algorithm performs 200 rounds of selection; each round consists of sampling three individuals from the offspring and selecting the one with the highest predicted promoter activity based on our K562 model (W). The 200 selected individuals compose a new generation that will start a new iteration of the algorithm. We ran the GA in five independent instances, each of which iterated throughout 500 generations. **b,** Gain in predicated promoter activity in one of the GA instances. Red curve indicates the mean predicted promoter activity of the population throughout the iterations; shading marks the full range of activities. **c-e,** Experimental validation of the activity of synthetic promoters. **c,** As reference, we first selected 27 of the strongest promoters in the human genome according to the input MPRA data^18^ (blue dots; violin plot shows activity distribution of all 30,607 promoters). From each of these 27 promoters we then selected a 230 bp fragment with the strongest PARM-predicted activity. **d,** PARM-predicted activity of the 20 synthetic promoters (red) compared to the predicted activity of the 27 native human promoter fragments (blue). **e,** Measured promoter activity of the same fragments as in **d**. The synthetic fragments are significantly more active than the strongest human promoter fragments (P = 0.00941, Student’s t-test). **f,** TF motif counts in the 20 synthetic sequences, detected by FIMO scanning^20^ with motifs in Hocomoco v1v11^21^. For each motif family, we only annotated the most frequent motif in the heatmap. Row colours indicate the replicates of five independent GA runs (4 sequences per run were selected). Sequences from the same runs tend to cluster as they share the same motifs. Hierarchical clustering was performed using complete linkage and euclidean distance as metric.

#### PARM has learned functional elements and regulatory grammar

Notably, the synthetic promoters are dissimilar to any human genome sequence (based on a BLAST^22^ search; see Methods), indicating that PARM is not simply reproducing sequences observed within human promoters. Together, these results demonstrate that PARM not only reliably recapitulates the activity of naturally occurring promoters, but also can produce highly active promoters *de novo*, by combining binding motifs of TFs that it has learned to be active in the tested cell type. This suggests that the model has learned biologically relevant aspects of the regulatory grammar of human promoters.

### Promoter-focused MPRA library allows scaling to multiple cell types

#### Design of a promoter-focused MPRA library

Next, we sought to further broaden this approach to a wider range of cell types. However, MPRA experiments with full-genome libraries as described above are technically challenging and require very large numbers of cells due to the high complexity of these libraries^18^. This substantially limits the scalability. Because we used only data from promoter-overlapping fragments to train PARM, we reasoned that an MPRA library containing only such promoter fragments would be sufficient. We therefore employed a capture-based strategy to create a new library that was highly enriched (91.3%) for promoter-overlapping fragments from a human genome (**Figure S2a-d**). This library consisted of ∼8.5 million sufficiently represented unique fragments, which is ∼100-fold less than in the combined full-genome libraries^18^, but with an average 302-fold coverage of all human TSSs (**Figure S2c**) and including a broad diversity of fragment sizes (**Figure S2b**).

#### Focused library yields a robust and reproducible model

We first assessed the performance of the promoter-focused library in K562 cells. The original whole-genome MPRA dataset was obtained by transfecting a total of ∼100 million cells for each of the eight libraries and for each replicate experiment^18^ while for the focused library we used only 10 million cells (typically one 10 cm dish) per replicate experiment. Strikingly, the overall predictive power of the model was highly similar between the two approaches (R = 0.91 versus 0.90, **Figure S3a, b**). Moreover, the PARM model obtained with the focused library was highly consistent when trained on independent biological replicates (average R = 0.92). This reproducibility was substantially higher than the reproducibility of the input data (average R = 0.82; **Figure S3c**), indicating that PARM has a potent de-noising effect and effectively focuses on the relevant signals in the data. The training of PARM for a single cell type requires only five hours using 12 cores, 10G of RAM and one RTX6000 GPU. Thus, both experimentally and computationally the promoter-focused design represents a highly economic strategy.

#### Expansion to other human cell types

We then generated additional promoter-focused MPRA datasets and PARM models from colon carcinoma (HCT116), breast cancer (MCF7), and prostate cancer (LNCaP) cells (**Figure S3d-f**), again requiring only 10 million cells for each replicate experiment. The estimates of predictive accuracy in these cells were lower (R = 0.65 to 0.81), but this may be attributed to the higher noise level in the input data (average R between replicates ranging from 0.42 for LNCaP to 0.8 for HCT116; **Fig S2e**), which puts upper limits to the validation estimates. These combined experimental and computational efforts yielded predictive models of promoter activity in five human cell types of highly diverse lineages.

### Identification of critical promoter sequences

#### PARM identifies critical sequences in TERT promoter

The thus established models enabled us to conduct for each promoter and in each cell type an in silico saturating mutagenesis that predicts the impact of individual nucleotide substitutions on the activity of the promoter^6,8,23^. To assess the utility of these maps we first examined the human TERT promoter, which has been extensively characterised because of its importance in cancer^24,25^. PARM correctly predicted that the mutations C250T and C228T, which occur in ∼25% of human cancers and result in increased TERT expression^26,27^ (**Figure 3a**, black asterisks). Interestingly, the model also predicted elevated promoter activity for several other TERT promoter mutations that are found at low frequencies in human tumours (**Figure 3b**). This suggests that these rare mutations may in fact contribute to oncogenesis in some cancers. Furthermore, PARM predicted five patches of 6-8 adjacent nucleotides that contribute to the activity of the promoter (**Figure 3a**). This is largely in accordance with a previous experimental mutagenesis study that found four of these sites to boost activity of the TERT promoter (**Figure 3a**, black bars)^28^. Thus, even though PARM was never trained on mutations in the TERT promoter, it recapitulated several known aspects of its regulatory code and made additional plausible predictions.

**Figure 3.**
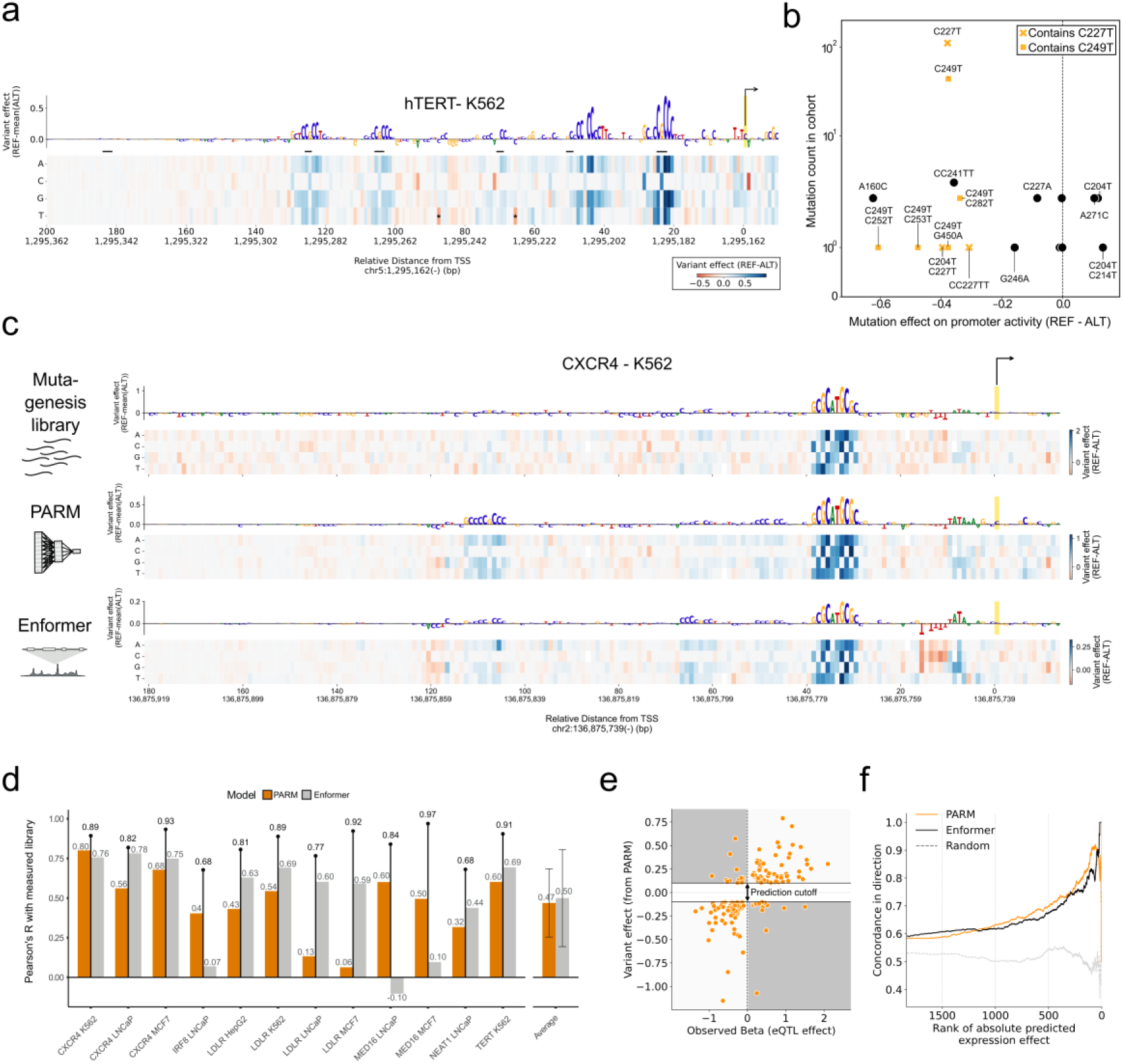
Single-nucleotide functional predictions by PARM. **a,** *In silico* mutagenesis analysis of the human TERT promoter in K562 cells. Heatmap shows the increases (red) and decreases (blue) in promoter activity predicted by PARM for all possible single-nucleotide substitutions throughout the promoter. Letter heights above the heatmap shows importance scores for every position, i.e. the average impact of the three possible substitutions on gene expression, relative to the reference sequence. A positive importance score means that the wild-type nucleotide positively contributes to promoter activity; a negative score implies that mutations enhance promoter activity. Horizontal bars mark nucleotides that were previously reported^28^ to be important for full promoter activity (black) or not important (grey). **b,** Most mutations found in the TERT promoter in the ICGC cancer cohort (n = 6241 distinct donor)^29^, including very rare ones, are predicted by PARM to lead to increased promoter activity. REF, reference genome allele; ALT, mutated allele. **c,** Effects of single-nucleotide substitutions across the CXCR4 promoter in K562 as measured by MPRA (top) and predicted by PARM (middle) or by *Enformer* (bottom). TSSs in **a, c** are marked by yellow bars; the direction of transcription is indicated by arrows. **d,** Pearson correlations between effects of single-nucleotide substitutions as measured by MPRA and effects as predicted by PARM (blue bars) and *Enformer* (grey bars), in eight different promoters tested in four different cell lines as indicated. Lollipops indicate Pearson’s correlations between independent replicate MPRA experiments. **e,** Concordance of the measured effects of a set of SNPs in human promoters according to fine-mapped eQTL data in whole blood^30^ with the predicted effects of these SNPs in K562 cells according to PARM. This panel shows the 125 SNPs with the strongest PARM-predicted effect sizes. Concordance (the proportion of SNPs in green areas) is 81.7%. **f,** Concordance with eQTL data for various effect size cutoffs, comparing predictions by PARM, *Enformer* and randomly sampled predictions taken from a uniform distribution.

#### Validation by experimental mutagenesis

To further test the PARM predictions experimentally, we created a synthetic MPRA library of a set of human promoters in which we systematically substituted every nucleotide with the three alternative nucleotides. We then measured the regulatory impact of each mutation in our own cultures of K562, HepG2, MCF7 and LNCaP cells (**Figure 3c,d**; see Methods). For 12 of these promoter/cell line combinations, we obtained measurements that passed stringent quality criteria (see Methods). Across these 12 tests, the Pearson correlation coefficient R between measured and predicted mutation effects was 0.49 ± 0.22 (**Figure 3c, d**). Replicates of the measured data correlated with R = 0.84 ± 0.09, indicating that perfect predictions should not be expected. We compared this performance to that of *Enformer,* a state-of-the-art DL model of gene activity that was trained on thousands of epigenome and transcriptome maps^8^. *Enformer* predicted the measured mutagenesis data with R = 0.50 ± 0.30 (**Figure 3c, d**), *i.e.*, with similar accuracy, but it requires about ∼1,000 times more computational time than our CNN model for each mutagenesis scan (see Methods).

#### PARM can predict cis-eQTLs

Finally, we tested the ability of PARM to predict the effects of promoter sequence variants on gene expression in human tissue. For a set of 1,836 fine-mapped cis-eQTLs (variants significantly associated with a change in gene expression, within ±1 Mb window around TSS) from GTEx v8 in blood samples^30^ we determined whether the allele that associated with an increase in gene expression is also predicted to have a higher PARM effect, compared to the allele that decreases expression. For 81.7% of the cis-eQTLs where PARM predicted a substantial difference between the two alleles (N = 125 SNPs with an absolute predicted effect above 0.1), we observed directional concordant effects (**Figure 3e**). At various effect size cutoffs for the top 1000 SNPs, PARM performed similarly to *Enformer* in this prediction task (**Figure 3f**). Thus, PARM can predict eQTL effects reasonably well, particularly considering that K562 cells, which are chronic myeloid leukaemia cells, are not a perfect match for whole blood.

### Inference of functional TF binding sites at promoters genome-wide

Encouraged by these results, and taking advantage of the highly efficient calculations, we applied the in silico mutagenesis analysis to 30,607 well-annotated human promoters (available in External Data 1). This typically identified one or more patches of 4-10 adjacent nucleotides that consistently were predicted to either enhance or reduce the activity of any particular promoter (**Figure 3a, 3c**). To identify the putative responsible TFs, we systematically compared these patterns to a database of known binding specificities of all TFs^21^. If the binding specificity of a particular TF correlated with |R| >*=0.75* at a given position, we assigned this TF to that patch as a putative regulator (**Figure 4a**; see Methods). A random permutation approach indicated that this cutoff for R yields a false-positive discovery rate <5% for virtually all TFs (**Figure S5**). As will be illustrated below, we emphasise that these maps do not simply describe putative TF binding sites based on DNA sequence alone, but rather *functional* binding sites where the TFs are predicted to *impact* promoter activity in the cell type for which the model was derived. For simplicity, we will refer to these sites as regulatory sites (RSs).

**Figure 4.**
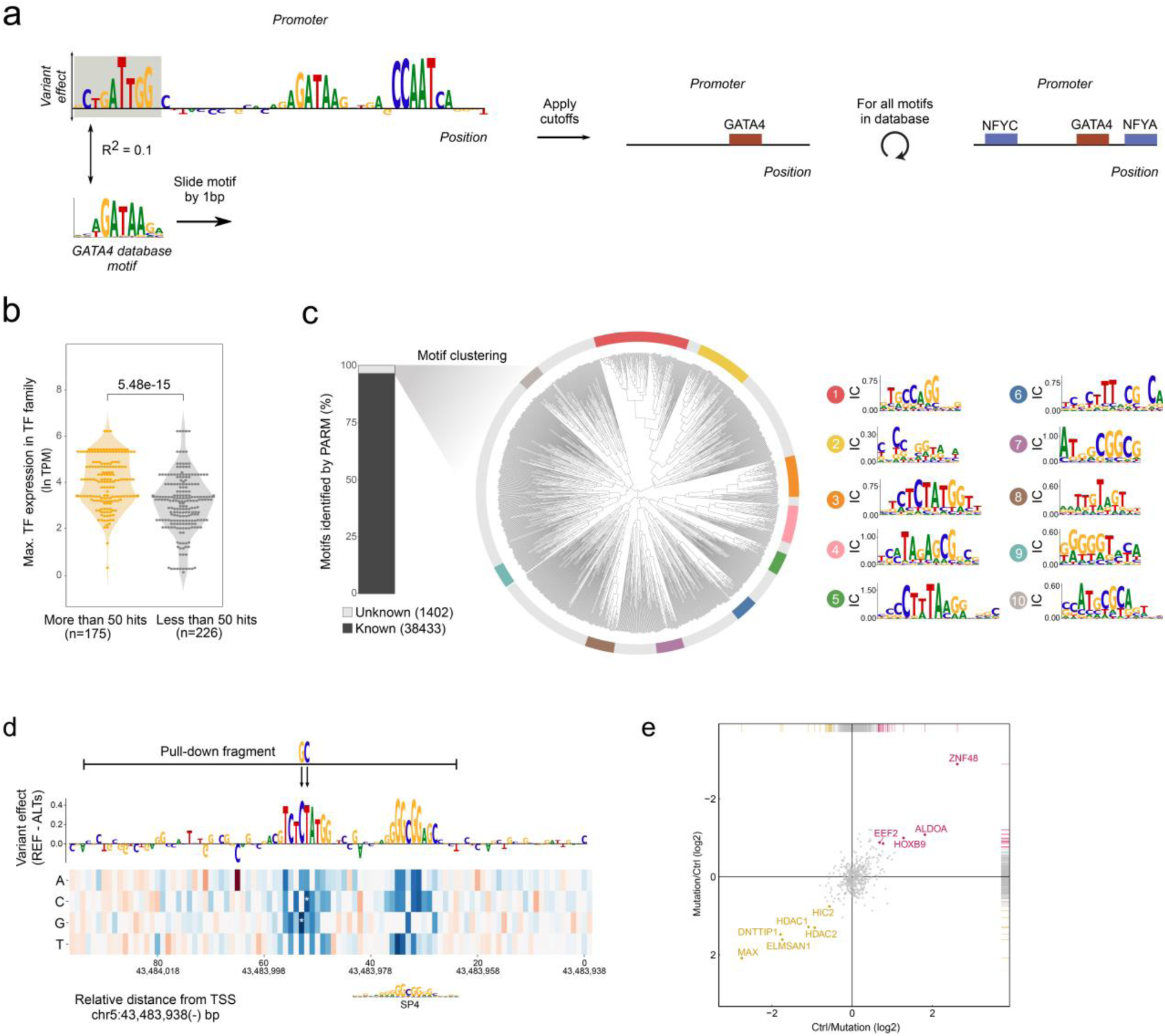
Activity inference of TF motifs in 30,607 promoters. **a**, Schematic representation of motif scanning and RS identification. Each TF motif in Hocomoco v11^21^ is scanned against the importance scores with a step size of 1 across the entire promoter. An RS is assigned if the TF motif is highly similar to the importance score (|R| > 0.75; see Methods). This process is repeated for all TF motifs and for all promoters. **b,** TFs with motifs frequently identified in RSs across all promoters (> 50 times) are more highly expressed than TFs with motifs that are rarely identified in RSs (<50 times). TF expression estimates are based on mRNA-seq data from Human Protein Atlas version 23.0^31^. If multiple TFs match a motif, then only the most highly expressed TF is considered. Results are shown for K562 cells. P-value is according to Mann–Whitney U test. **c,** In a separate approach, all promoters were screened for regions consisting of at least five consecutive nucleotides with high absolute variant effect scores. 96.5% of these regions corresponded to known TF motifs (see Methods). Hierarchical clustering of the remaining 3.5% of regions yielded 10 clusters of at least 30 regions with similar importance score patterns (coloured sections of circular cluster diagram). The consensus patterns for each of these clusters are shown using the Information Content (IC). **d,** *In silico* mutagenesis of TMEM199 promoter computed in a region of −250 bp to 50bp relative to TSS, showing only the relevant region. The TCTCATGG motif does not match any motifs in Hocomoco v11^21^ or Jaspar2024^32^ database. A pull-down assay was conducted using a 70bp DNA fragment marked by the horizontal bar. The reference sequence is compared to a sequence with the putative RS mutated at positions −52bp and −53bp relative to the TSS. **e,** Pull-down results comparing the proteins binding to the wild-type TMEM199 promoter and the mutated version as indicated in **d.** The experiment was performed once with heavy labeling of proteins bound to the wild-type DNA fragment (Ctrl) and light labeling of proteins bound to the mutated DNA fragment (*x-*axis), and once with reverse labeling orientation (*y-*axis). Proteins highlighted in the upper-right quadrant bind poorly to the mutated DNA fragment. See **Figure S7** for additional experiments.

#### Assignment of TFs to most promoters; PARM primarily captures expressed TFs

Most of the TFs that were identified by PARM as contributors to promoter activity were indeed expressed in the respective cell type according to mRNA-seq data. In the majority of cases where the model assigned regulatory activity to a TF that was actually not expressed, the cognate motif was highly similar to that of an expressed TF; when this was taken into account, virtually all detected motifs could be linked to an expressed TF (**Figure 4b**). To avoid such erroneous assignments, we repeated the motif matching after removing all non-expressed TFs. In K562 cells this resulted in 20,543 promoters with at least one RS (**Figure S6a**). We note that many RSs matched multiple TFs that have highly similar binding specificities and thus were annotated more than once. As may be expected, the remaining 10,064 promoters without any predicted RS were generally much less active in the MPRA as well as endogenously (**Figure S6b**).

#### The majority of RSs match known TF motifs

Next, we asked whether there may be additional RS in human promoters that do not match any currently known TF binding motif. For this, we systematically scanned our maps of all promoters for patches of at least 5 adjacent nucleotides that consistently were predicted to either enhance or reduce the activity of a promoter (see Methods) but did not match any known TF binding motif at the cutoff that we applied. In K562 cells this yielded 1,402 of such unannotated RSs, whereas 38,434 RS matched at least one known TF motif (**Figure 4c**, left panel). Thus, PARM converged strongly on known TF biology, even though it was never provided with TF motif information. Unsupervised clustering of the 1,402 un-annotated RS sequences (see Methods) resulted in the identification of 10 clusters that contained at least 30 similar RSs each (**Figure 4c**, middle and right panels).

#### Cluster 3 motif is bound by ZNF48

We further investigated motif cluster 3 with 57 hits in the human promoters, with consensus sequence TCTCTATGGT. To identify the putative TF that binds this motif, we incubated DNA fragments containing this motif, or containing mutated versions of the motif, with K562 nuclear extracts and identified the bound proteins^33^ by stable isotope labeling-based (**Figure 4d-e; S7b-f**) and label-free based (**Figure S7g-h**) quantitative mass-spectrometry. This identified ZNF48 as the prominent and most consistent candidate TF interacting with this motif across all biological replicate experiments. Although the motif of ZNF48 is not listed in current databases, a TCTCT motif was inferred for this TF based on chromatin immunoprecipitation data^34^, which is in agreement with our proteomics data. This result illustrates the sensitivity of PARM, as it can identify functional TF motifs that are relatively infrequent and poorly annotated.

### Activating, repressive and cell type-specific TF functions

#### Activating, repressive and dual-function TFs

RSs for which in silico mutations predict loss of activity are likely to be bound by an activating TF, while RSs for which mutations consistently lead to a gain of activity are likely to be bound by a repressive TF (**Figure 5a-c**). Strikingly, some RSs matching the same TF motif showed opposite effects depending on the promoter. For example, NRF1 is predicted to activate the *POMGNT2* promoter (**Figure 5a**) but repress the AC004834.1 promoter (**Figure 5c**), even though the RS in both cases is located immediately upstream of the TSS. For a global view of this dichotomy, we first surveyed the abundance of motif matches for each TF, separately for activating and repressive RSs (**Figure 5d**). This identified several TFs predicted to function prominently as repressors, such as ZEB1, SNAI1 and ZBTB7A. All of these have long been known to be repressors^35–37^. Other TFs, such as several members of the KLF, ELK, ELF and SP families are identified primarily as activators, in agreement with earlier studies^38–40^. Interestingly, the model also predicts that many TFs have both repressive and activating functions. This includes known dual-function TFs such as IKZF1^41^ and MYC^42^, yet the model predicts that many other TFs can act both as activators and as a repressor, such as NRF1 (**Figure 5a-c**). Below we will further characterise possible determinants of this functional dichotomy.

**Figure 5.**
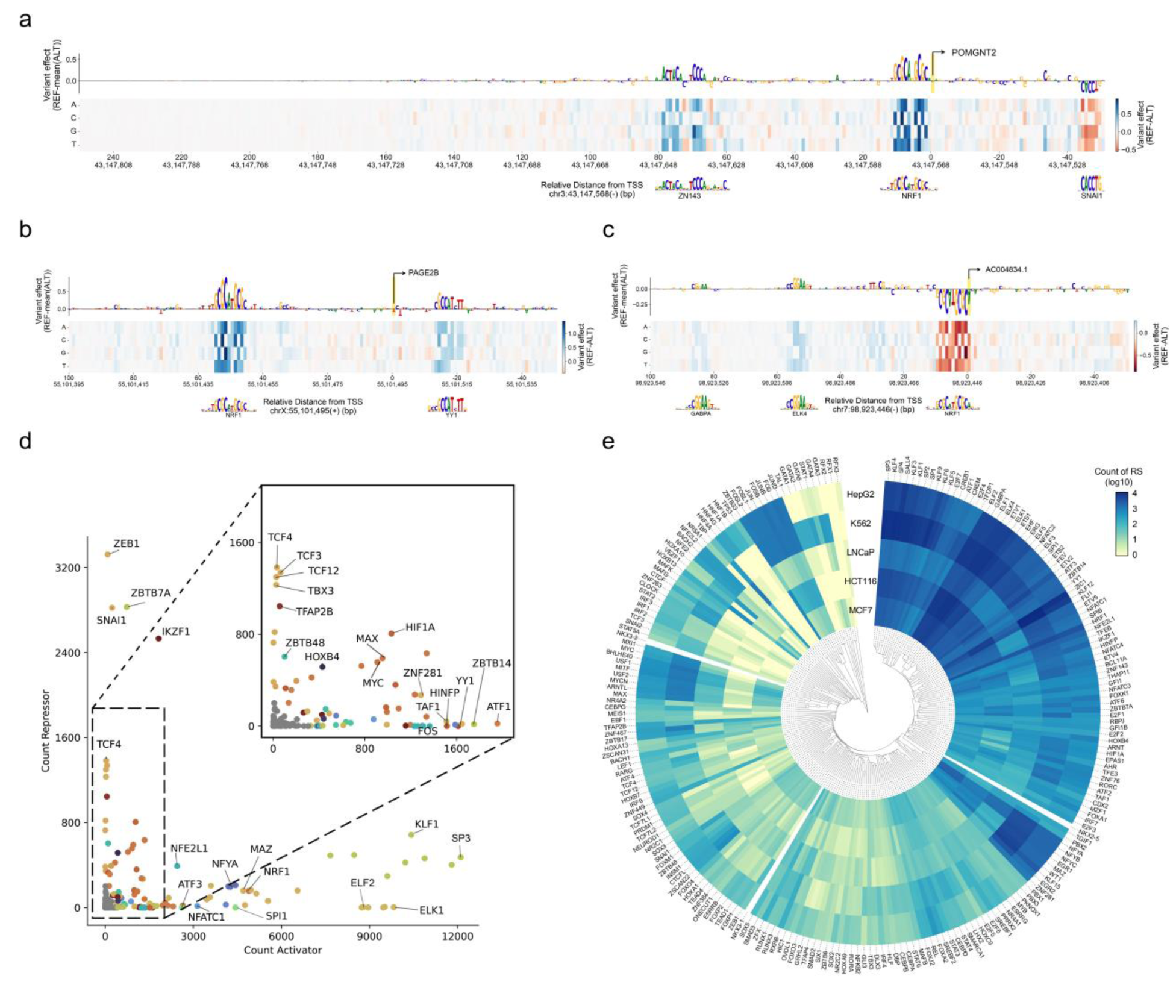
Activating and repressive functionality, and cell type specificity of TF motifs. **a-c,** Examples of promoters in K562 cells with activating and repressive RSs, with best-matching TF motifs. Note that NRF1 is predicted to act both as an activator or repressor, depending on the context. Yellow bar: TSS; arrow: direction of transcription. **d,** Total counts of repressive and activating RSs throughout all 30,607 promoters in K562 cells, stratified by overlap with individual TF motifs. TF motifs from the same family are shown in the same colour (see Methods). Note that many TF motifs are predicted to be both repressive and activating. **e,** Clustered heatmap showing counts of activating RSs overlapping with each TF motif, for five different cell types. Hierarchical clustering was performed using complete linkage and Euclidean distance as metric.

#### Cell type-specific and shared regulators in human promoters

It is generally thought that cell type-specific regulation of gene expression is primarily controlled at the level of distal enhancers, rather than at promoters. In agreement with this notion, our models of TF activity at promoters show strong similarities between all tested cell types, both for activating and repressive activities (**Figure 5e**). Nevertheless, some TFs exhibit strong differences between cell types. For example, the Hepatocyte Nuclear Factor (HNF) associated RS are restricted to HepG2 cells, while GATA factors, which are highly active in blood-related cells^43^ are most prominently detected by PARM in K562 cells (**Figure 5e**).

### Preferred positioning of functional TF motifs

#### Global positional preference of TF activity

Having identified putative RSs and their associated TF motifs throughout 30,607 human promoters, we then investigated whether TFs may have specific positional preferences relative to the TSS for their regulatory activity. All RSs aggregated over all promoters showed preferential positioning around −120…+10 bp, with a peak at −50 bp (**Figure 6a**). This is in close accordance with previous estimates based on linear regression of promoter MPRA data^1,44^, indicating that this distribution is not a result of a bias of the CNN model towards TSS-proximal elements, but rather reflects a biological pattern. We compared this result to standard TF motif scanning based on sequence alone^20^. This showed a much flatter distribution and identified about 10 times more motif matches than PARM (**Figure 6a**). This suggests that the majority of TF motifs in promoters are not functional, possibly because they lack proper sequence context (see below), although we cannot rule out that PARM fails to detect activity for a subset.

**Figure 6.**
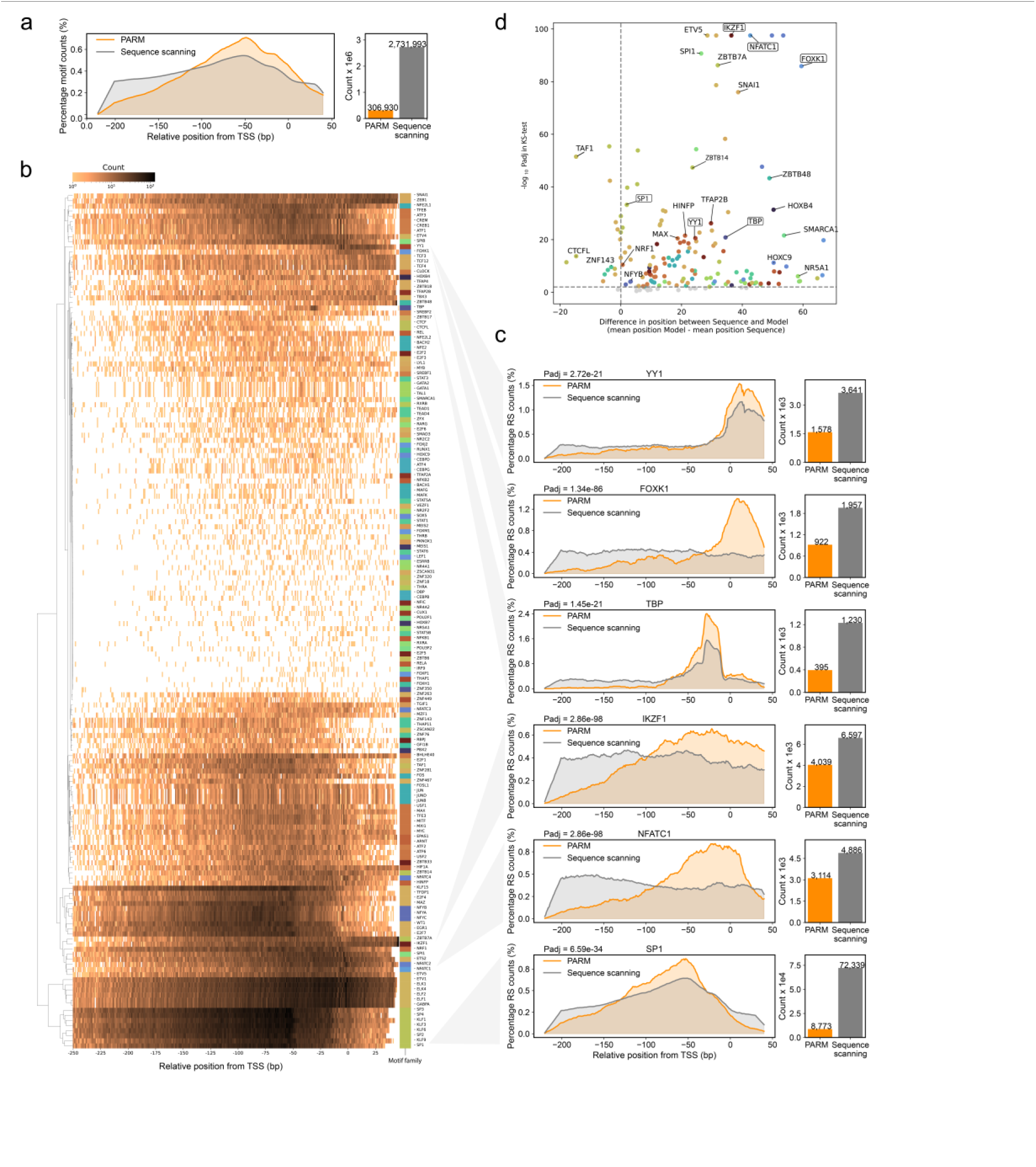
Distinct preferential positioning of functional TF motifs in K562 cells. **a,** (left panel) Distribution of all RSs matching a TF motif (orange), compared to the distribution of all combined TF motifs as identified by plain sequence scanning using FIMO^20^ (grey), aggregated across all 30,607 promoters; (right panel) corresponding numbers of TF motif matches. **b,** Distribution of RSs relative to the TSS, stratified by matching TF motif. Hierarchical clustering was performed using complete linkage and Euclidean distance as metric. Colours next to TF names depict TF motif families. **c,** Detailed RS distributions for selected TF motifs (orange) compared to the distributions of the same motifs as detected by plain sequence scanning (grey). Curves in (a, c) were smoothed by linear convolution with a 20 bp sliding window. To test for the difference in distributions, P-values were calculated with the Kolmogorov–Smirnov (KS) test and adjusted with the Benjamini-Hochberg method (Padj). **d,** Positional biases of functional TF motifs; a positive value indicates that functional motifs identified by PARM tend to be downstream of corresponding motif incidences identified by FIMO. Padj calculated as in **c.** Boxed TF motif names are highlighted in **c**. Colours indicate motif families as in **b**.

#### Individual TFs have distinct regulatory positions

We then repeated this pan-promoter analysis for RSs belonging to individual TF motifs. Strikingly, this uncovered a diversity of patterns that differ substantially between TFs (**Figure 6b** and **Figure 6c**, orange curves). For example, YY1 and FOXK1 are most active just downstream of the TSS, while NFATC1 acts preferentially in the window −100 bp to +20 bp, and SP1 mostly acts further upstream, with a peak of about 50 bp upstream of the TSS. Reassuringly, RSs controlled by TBP are strongly enriched at position −30 bp, which is exactly where TBP is known to act^45^.

#### Sequence motifs poorly explain positional preferences

Importantly, the positional preferences of each TF are only partially explained by the location of their cognate motifs. The TBP binding motif (TATA box) is indeed prominent around −30 bp but it is also found further upstream, whereas the model predicts that it is rarely active at those more distal positions (**Figure 6c**, grey versus orange curve). For most other TFs, the cognate motifs are much more homogeneously distributed across the entire promoter regions than the RSs identified by PARM, which can be considered *functional* motifs. We find such differences between the positioning of the functional motifs versus all motifs detected by plain sequence scanning to be statistically significant for 150 out of 247 TFs (**Figure 6d**). Thus, PARM provides functional information that cannot be obtained by simple scoring of TF motif occurrences in promoter sequences.

### Positional preferences differ for activating and repressive functions

#### Repressive RSs tend to be near or downstream of the TSS

We then wondered whether the positional preference of RSs may differ between activating and repressive RSs. Strikingly, this is indeed the case: overall, the distribution of activating RSs has its mode around position −50 bp, while repressive RSs are most abundant closer to the TSS, roughly between positions −25 and +50 bp (**Figure 7a**). Notably, this spatial separation of function is also true for dual-function TFs. For the majority of these TFs we found that activating RSs are more upstream of the TSS than repressive RSs (**Figure 7b**), even though the sequence motifs are not detectably different. For example, for RSs that bind IKZF1, repressive sites show a marked enrichment downstream of the TSS, while activating sites are almost exclusively upstream (**Figure 7c**). A similar strong spatial separation is seen for BHLHE40; for several other dual-function TFs the repressive RSs are positioned mostly upstream, yet more closely to the TSS than the respective activating RSs (**Figure 7c**).

**Figure 7:**
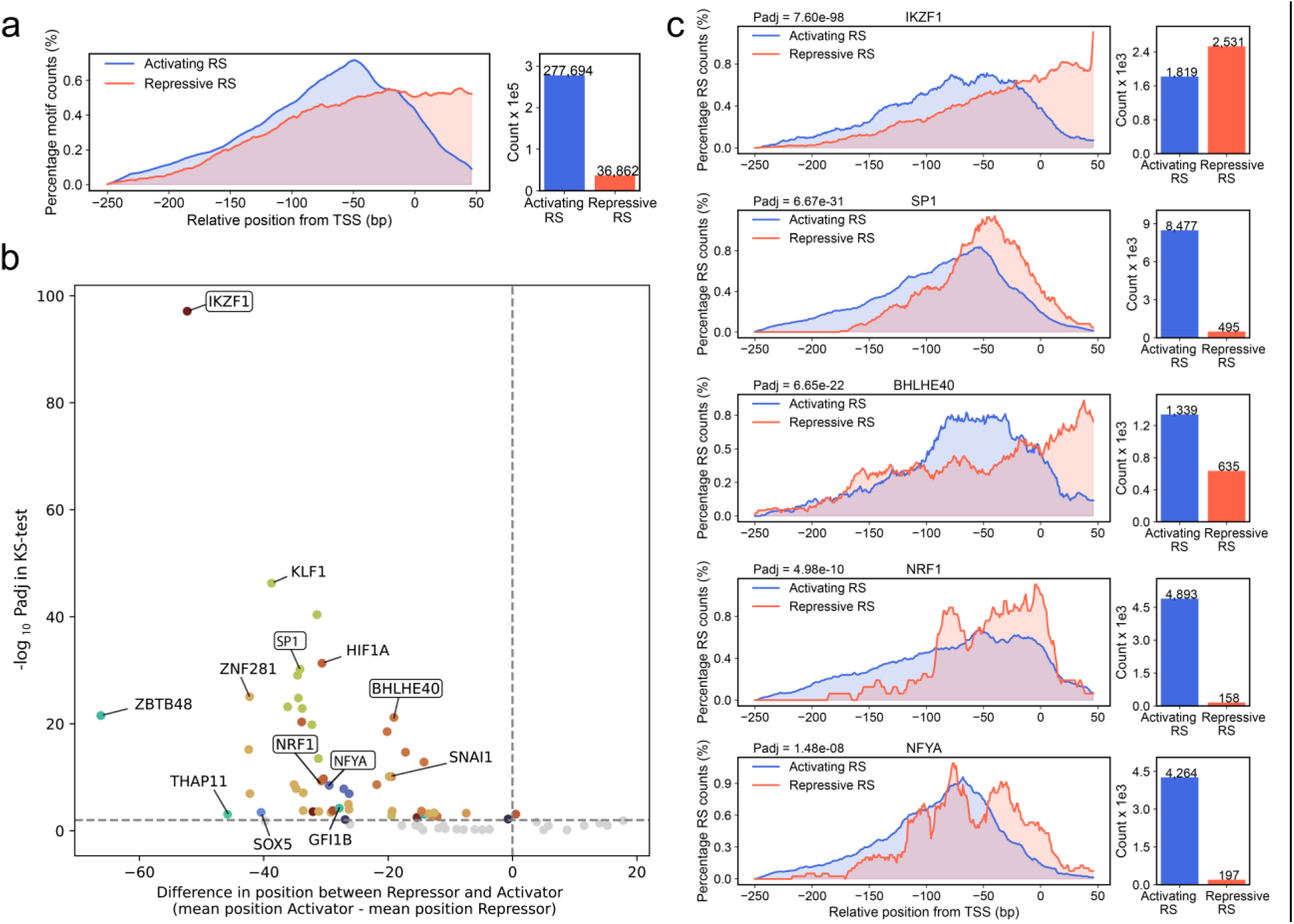
Distinct positional preferences of activating and repressive TF motifs. **a,** (left panel) Distribution of all RSs identified by PARM in 30,607 promoters as activating (blue) or repressive (red). (right panel) Activating RSs are ∼7.5 times more common than repressive RSs (*cf.* **Figure 5d**). **b,** Majority of bifunctional TFs show a systematic downstream shift of repressive RS, compared to activating RS matching the same motif. P-values were calculated with the Kolmogorov–Smirnov (KS) test and adjusted for multiple testing with the Benjamini-Hochberg method (P_adj_). Boxed TF motifs are highlighted in **c.** Colours indicate motif families. **c,** Detailed RS distributions for selected bifunctional TF motifs, comparing activating (blue) and repressive (red) RSs. Curves in **a, c** were smoothed by linear convolution with a 20 bp sliding window. Results in **a-c** are from K562 cells.

#### Possible mechanism for repression

We propose that TFs that bind to repressive RSs that are close to the TSS may simply act by steric hindrance of pre-initiation complex assembly, while repressive RSs that are located downstream of the TSS may act as (partial) roadblocks for transcription elongation. Transcriptional roadblock activity of TFs has been extensively documented in prokaryotes (reviewed in ^46^). It has also been observed for a few TFs in yeast^47–50^ and for CTCF in human cells^51,52^, but it has not been systematically linked to eukaryotic promoter regulation.

### Combinatorial and positional grammar revealed by *in silico ins*ertion of TF motifs

#### Motif insertion scans uncover complex grammar

To further explore the impact of TF motif positioning, we computationally inserted individual TF motifs systematically throughout native promoter sequences and used PARM to predict the impact on transcriptional output. This revealed a high diversity of position-dependent effects. For example, the NRF1 motif could activate or repress the *FCF1* promoter in a strongly position-dependent manner (**Figure 8a**). When placed upstream of the TSS in silico, it showed a predicted activating effect that gradually strengthened with decreasing distance to the TSS. In contrast, when it was inserted at or downstream of the TSS, it showed repressive activity. Interestingly, PARM indicates that this repression involves the muting of the activating RS that is located 5’ of the NRF1 motif insertions. None of these effects were observed when these simulations were repeated with two different mutated NRF1 motifs (**Figure S8a**), indicating that the effects are not simply due to altered spacing of RSs.

**Figure 8:**
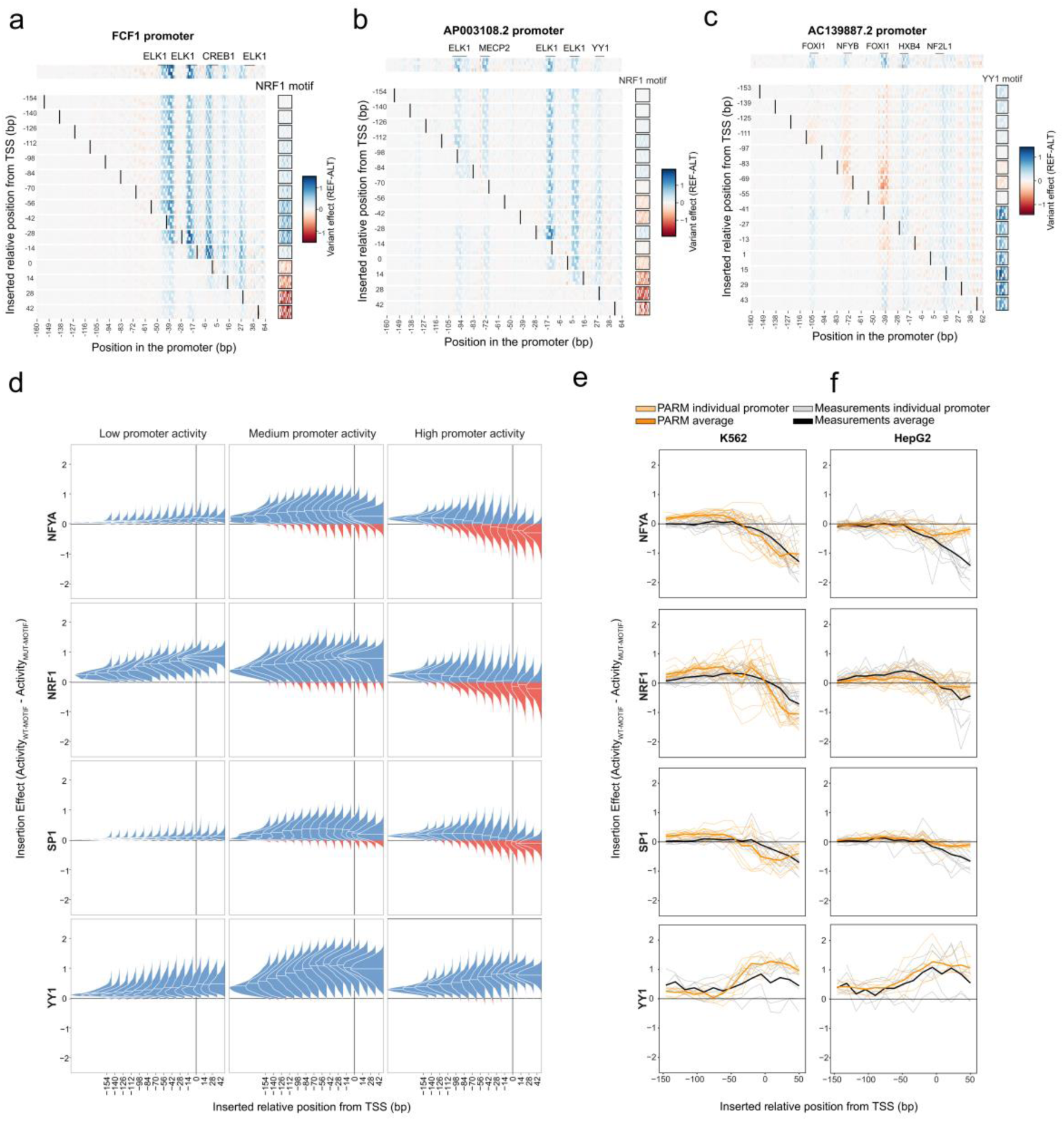
PARM reveals the positional grammar of TF motifs. **a-c**, *Insilico* TF motif insertion scans for three different promoters using the K562 PARM model. The upper row shows the mutagenesis matrix of the native promoter, with RSs and their best-matching TF motifs indicated. The rows below show the mutagenesis matrices after systematic insertion of a TF motif (size 12bp for YY1 and 14bp for NRF, SP1 and NFYA) at positions marked by vertical black bars. The activity of the motif itself is shown to the right of each matrix. **a,** insertion of the NRF1 motif in the *FCF1* promoter. **b,** Insertion of the active NRF1 in the *AP003108.2* promoter. **c**, insertion of the active YY1 motif in the *AC139887.2* promoter. See **Figure S8a-c** for the same analyses with mutated TF motifs, which generally do not have the same effects. **d,** Motif insertion scans with NFYA, NRF1, SP1 and YY1 motifs for all 30,607 human promoters. The y-axis shows the difference in the predicted activity of each promoter after the insertion of the intact TF motif (ScoreActive) and the average predicted activity of the same promoter after the insertion of corresponding mutated TF motifs (ScoreInactive). Results are split by wild-type promoter activity (High: highest quintile; Medium: middle three quintiles; Low: lowest quintile). Violin plots show score distributions across the tested promoter sets, with their bases marking the position of motif insertion relative to the TSS. Blue: promoter is activated by the motif insertion; red: promoter is repressed by motif insertion. **e-f,** Experimental insertion mutagenesis MPRA confirms positional effects. For each of the four TFs in **d**, sets of synthetic promoters of 230 bp long were subjected to systematic insertion mutagenesis with intact and mutated TF motifs. Plots show the effects of the insertions of intact motifs relative to matching mutated motifs on promoter activity when transfected into K562 cells (**e**) and HepG2 cells (**f**). Thin grey curves represent individual promoters measured by MPRA; thick black curves show corresponding averages. For reference, orange curves show predictions by PARM (thin: individual promoters; thick: average).

#### Example of motif interactions

The same NRF1 motif exhibited more complex behaviour when inserted throughout the *AP003108.2* promoter (**Figure 8b**). Like in the *FCF1* promoter, it initially showed a gradually increasing activating function when moved from position −154 to −84. But closer to the TSS it switched between repressive and activating functions, possibly by interacting with surrounding RSs. When placed at or downstream of the TSS the NRF1 motif acted again as a repressive element and muted several motifs located 5’ of the insertion, similar to the *FCF1* promoter. Again, these effects were not observed with a mutated NRF1 motif (**Figure S8b**).

#### YY1 motif shows different effects

A third example is the YY1 motif inserted throughout the *AC139887.2* promoter (**Figure 8c**). Again, depending on its precise position, the YY1 motif showed activating, repressive or neutral effects. In contrast to the NRF1 motif, however, it showed potent activation activity when inserted near or downstream of the TSS. None of these effects were observed with a mutated YY1 motif (**Figure S8c**). These examples illustrate that PARM does not predict promoter activity by simple addition or multiplication of individual motif activities, but instead captures a complex combinatorial and position-dependent grammar. seemed at odds with the global positional trends that we observed for RS in their natural positions. To investigate this, we selected four TF motifs of 12-14 bp and systematically inserted their sequence computationally throughout all 30,607 promoters, and calculated the predicted effect on transcriptional output, compared to mutated motifs inserted in the same positions. For a more detailed view, we grouped the promoters into three classes based on their predicted expression levels: the highest quintile, the combined three middle quintiles, and the lowest quintile (**Figure 8d**). The results of this computational insertion mutagenesis are largely in agreement with the positional effects observed for the respective motifs in their natural positions and provide additional insights into the functionality of each TF motif. For example, in the quintile of most active promoters, NRF1, SP1 and NFYA motifs showed the strongest activating effects around positions −150 to −50, and were predominantly repressive when inserted near or downstream of the TSS; while YY1 showed the opposite trend, with the highest activity near or downstream of the TSS (**Figure 8d**, right panels).

#### Some TF effects depend on promoter expression level

Surprisingly, in promoters in the lowest expression quintile, different patterns were visible (**Figure 8d**, left panels). Here, NRF1 was exclusively activating, particularly when located near the TSS. Furthermore, NFYA and SP1 motifs showed very little activity in this quintile, suggesting that these TFs can only activate promoters that have a certain baseline activity, presumably due to the presence of other activating TFs. Together, these results indicate that positional functional biases and complex motif–motif interactions constitute two global principles of promoters.

### Experimental data confirm positional effects

#### MPRA testing of sequences with insertions

To test the general validity of these predictions, we constructed a synthetic MPRA library in which we systematically inserted four different TF motifs and two mutated motifs for each (the latter predicted to be inert) at 15 different positions in 19 different promoters. We transfected this library into K562 and HepG2 cells and measured the effect of each motif insertion relative to the matching mutated controls. These experiments confirmed that the NFYA, NRF1 and SP1 motifs adopt a repressive mode when inserted near or downstream of the TSS, while the YY1 motif most potently activates transcription in these positions (**Figure 8e, f**). In HepG2 cells, these measured repressive effects of the former three motifs appeared to be more pronounced than PARM predicted, suggesting that the current HepG2 model underestimates the repressive impact of motifs near or downstream of the TSS. The activating role of YY1 at downstream positions is in agreement with a recent report^13^, but the underlying mechanism is still unclear.

## DISCUSSION

In this work, we created PARM, a detailed model of TF regulation across all human promoters and multiple cell types. To achieve this, we combined high-throughput reporter assays and efficient convolutional deep learning, resulting in highly accurate models that are scalable enough to perform exhaustive in silico mutagenesis across all human promoters. We moreover demonstrate that PARM can effectively be trained using data from a promoter capturing strategy, allowing model creation and application across many different cell types at minimal loss in model predictive accuracy. Together, these feats open up the possibility of using PARM to study the effect of sequence variation and promoter architecture at an unprecedented scale.

Several lines of evidence indicate that the model captured biologically relevant information, without any prior knowledge of TF biology. First, the vast majority of RSs that PARM identifies match known TF motifs. Second, among these RSs, PARM correctly annotated and distinguished between known repressive and activating TFs. Third, using cell-type-specific PARM models we uncovered several RSs with known cell-type-specific TF activities. Fourth, PARM correctly predicted the known biology of the TERT promoter, including the effect of well-established cancer-relevant mutations, and its predictions match eQTL data. Fifth, experimental nucleotide-resolution mutagenesis data can be predicted with reasonable accuracy, with correlations in line with competing models and strong RSs in the correct location. Sixth, when used in an in silico evolutionary optimization strategy, PARM can create strong promoter sequences, which exhibit experimentally validated stronger or equal transcriptional activity than the most active endogenous promoters. Apart from its potential utility in designing sequences with certain biological characteristics, this latter result demonstrates the ability of the model to extrapolate to regulatory sequences never observed in the training data.

Nevertheless, we emphasise that the model is likely to contain many imperfections. Most of all, it is a model of isolated promoters, in the absence of distal enhancers, insulators, epigenetic modifications and large-scale chromatin domains. It thus should be regarded as a reductionist model that provides insights into the basic properties of promoters. Interactions with distal elements may be better captured by long-range models such as *Enformer*, although this model also has limitations^53,54^ and is only trained on correlative epigenome datasets. Importantly, *Enformer* is computationally much more expensive and thus not suitable for in silico mutagenesis analyses of ∼30,000 promoter sequences, which we demonstrated here to offer a treasure trove of mechanistic insights and testable hypotheses. In the future, hybrid models that include both MPRA data and epigenome maps may be developed that combine the advantages of both strategies.

Sequence-based expression prediction models hold the potential to predict the effects of non-coding mutations and assist in fine-mapping of germline variants. However, as we demonstrate with PARM, lightweight, high-performance computational models – trainable on limited datasets – can surpass this goal. PARM facilitates in-depth studies of TF biology at a scale unattainable through traditional experiments. For instance, we demonstrate that it becomes possible to predict activating and repressive activities of TFs in every promoter, and to study the positional effects of functionally active TF binding. Using PARM, we determined that TF binding position relative to the TSS is a crucial factor in dictating activating or repressive effects. Moreover, by in silico integrating RSs in various locations in selected promoters, we were able to study positional dependencies of these RS integrations and their dependency on and interaction with surrounding RSs.

Collectively, these analyses highlight the remarkable ability of sequence-based deep learning models to deliver testable hypotheses about the intricate interplay between sequence variation and promoter architecture at an unmatched scale. When combined with rigorous experimental validation, this approach offers a powerful method to decipher the fundamental principles of human gene regulation. Given the relatively low computational and experimental costs of our approach, we envision that it will serve as a tool for prioritising hypotheses across diverse cell types and environmental conditions, thereby accelerating the elucidation of gene regulatory grammar.

## METHODS

### TSS selection

The selection of relevant TSSs was done as previously described^55^. Briefly, we selected GENCODE defined TSSs^56^, with the additional requirement that they are active in at least one cell type or tissue according to the FANTOM5 database^57^. This resulted in a curated set of 30,607 TSSs.

### Construction of MPRA libraries

#### Focused library (DNA Fragmentation & capture-based)

To generate the focused library, 100 µg DNA was isolated from a human cell line (HG02601) using the Isolate II Genomic DNA kit (Bioline*, BIO-52066*). The isolated DNA was then fragmented using dsDNA Fragmentase (*NEBNext, M0348L*) for 30 minutes and subsequently size-selected by gel-extraction for fragments sizes ranging from ∼200-400 bp. DNA was end-repaired using the End-IT DNA End-Repair Kit (Lucigen*, ER81050*) and subsequently A-tailed using Klenow Fragment (3’ -> 5’ Exo-; *NEB, M0212M*). For cloning purposes, two custom 31 bp dsDNA adapters (oNK46, oNK47) containing a T-overhang for the 5’ and 3’ ends of the fragments were ligated to the fragments using TA-ligation with the Quick Ligation Kit (NEB*, M2200L*; see **Supplementary Table 1** for oligonucleotide sequences). These adapters contain overlaps with the 3’ and 5’ ends of the linear barcoded p101 vector (see^18^) to allow Gibson assembly of the fragments after hybridization. PCR amplification was performed using 8 cycles with primers oNK51 and oNK52 and Equinox polymerase (Twist, *#104176*). To capture the promoter region of the TSS, we selected 30,607 TSSs and their - 300 to +100 bp window and ordered a *high stringency* hybridization capture library from TWIST for these custom regions consisting of 127575 probes. To prevent nonspecific binding of probes to our fragments, we designed custom blockers complementary to the custom Gibson adapters (oNK57, oNK58). Fragments were captured using the custom hybridization panel according to manufacturer protocols. Briefly, 1 µg of fragments were hybridised to the custom hybridization panel for 16 hours in presence of specific (custom designed) and nonspecific (Cot-1 DNA) blocker solution. Subsequently, hybridised fragments were enriched using streptavidin binding and amplified using primers oNK51 and oNK52 using PCR for 9 cycles. Captured fragments were purified and cloned into the linear barcoded p101 vector with Gibson assembly using the HiFi DNA assembly Master Mix (NEB*, E2621L*) for 60 minutes at 50C. The Gibson assembly mix was then purified and subsequently transformed into MegaX DH10B ultracompetent bacteria (Thermo*, C640003*) according to manufacturer’s protocols. Transformed bacteria were grown in LB overnight and the plasmid library was isolated using the Gigaprep Isolation kit (Thermo*, K210009XP*).

#### Oligo-based libraries

All three synthetic libraries (synthetic promoters, in silico mutagenesis, and motif insertions) were generated using synthetic oligos that were ordered from Twist. Detailed design and count tables can be found in **Supplementary Data 1** for the synthetic promoter library, **Supplementary Data 2** for the in silico mutagenesis library and **Supplementary Data 3** for the motif insertion library. Oligo library was ordered from Twist and amplified using KAPA HiFi Hotstart Readymix (Roche*, KK2601*) for 14 cycles using oNK69 and oNK71. Library was bead purified, End-repaired using End-IT DNA End-Repair kit (Lucigen*, ER81050*), subsequently bead purified and digested with EcoRI-HF (*NEB, R3101S)* and NheI-HF (NEB*, R3131S*) for 1h at 37 C. Fragments were ligated in NheI-HF/EcoRI-HF double-digested vector (an adaptation of the p101 vector, see^58^) using T4 DNA ligase (Roche*, 10799009001)*. Ligation mix was purified and transformed as described above

### Cell culture and transfection

K562 was cultured in IMDM (for transfections) or RPMI1640 (lysate preparations), HepG2 and MCF7 in DMEM, HCT116 in McCoy5a, LNCaP in RPMI1640. All media were supplemented with 10% FBS (Gibco) and 1% penicillin-streptomycin (Thermo). All cells were transfected as described previously^18^, except HCT116, LNCaP and MCF7 were transfected using programs D-032, T-009 and P-020. For each replicate of the focused library 10 million cells were used. For each replicate of any oligo-based library, 5 million cells were used. All cells in culture were regularly tested for mycoplasma.

### Library sequencing

Library characterization (barcode-to-fragment association) the focused library was done by iPCR as described previously^18^. Library input (pDNA) was also generated as described previously, except here NcoI-HF was used to digest the plasmid library.

### RNA sample processing

RNA was isolated using Trisol (*Thermo*, #15596018) and divided into 10 µl reactions containing 2.5 µg RNA per reaction. Per biological replicate 5-10 reactions were done in parallel. To each reaction, we added 10 units of DNaseI for 60 minutes (*Roche,* #04716728001) which was subsequently inactivated by addition of 1 µl 25mM EDTA and incubation at 70°C for 10 minutes. Then, cDNA was generated by adding 1 µl of gene specific primer targeting the GFP ORF (oNK17) and 1 µl of dNTP (10mM each) and incubated for 5 min at 65°C. This reaction was continued by adding 4 µl or RT buffer, 20 units of RNase inhibitor (*Thermo*, EO0381), 200 units of Maxima reverse transcriptase (*Thermo*, EP0743) and 2.5 µl of water for 60 minutes at 50°C, and subsequently heat-inactivated at 85°C for 5 minutes. Each reaction was PCR-amplified with MyTaq Red Mix (Bioline, BIO-25043) using (index variants of) MMA219 and oNK28. PCR reactions were purified using bead-purification and subjected to single read sequencing on Illumina NextSeq or NovaSeq 6000 platforms.

### Read processing

Sequencing reads were processed as described previously^18^. This resulted in a table with read counts per individual fragment for both cDNA (fragment activity) and either iPCR or pDNA (input, normalisation) Then, we normalised the read counts as follows:

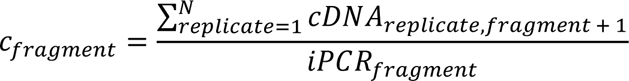

and defined Promoter activity (PA) as follows:

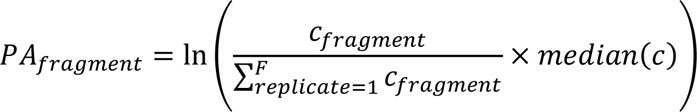

where *cDNA*_*replicate*,*fragments*_is the count for a specific fragment in a replicate, *iPCR*_*fragment*_ is the abundance of each fragment in the plasmid pool, *N* is the total number of replicates, and F the total number of fragments.

### Model input

PARM takes as input the genomic sequence of a single fragment from the library and predicts the associated promoter activity. This sequence is represented as a matrix of shape (600, 4) representing a one-hot-encoded DNA sequence: A = [1, 0, 0, 0], C = [0, 1, 0, 0], G = [0, 0, 1, 0], T = [0, 0, 0, 1], N = [0, 0, 0, 0], of length 600 bp. Since MPRA fragment lengths vary from 88 bp to 600 bp, we padded sequences shorter than 600 bp with N until reaching the maximum length. The number of N bases padded on the right and left sides of the fragment sequence was randomised for each fragment following a uniform distribution to avoid potential biases towards positions in the middle of the matrix. Additionally, to account for the creation of motifs on the edges of the fragments, we added adaptor sequences from the backbone plasmid. On both edges of each fragment, we included seven nucleotide sequences corresponding to the plasmid backbone. Specifically, for the genome-wide library, these sequences were CAGTGAT and ACGACTG, and for the focused library, they were CAGTGAT and CACGACG.

### Deep-learning model architecture

The PARM architecture, adapted from the first layers of Enformer^8^, consists of a CNN with self-attention pooling layers (**Figure S1**). Models were constructed individually for each cell type. The first layer comprises a 1D convolution layer of kernel size 7 and filter size 125, followed by a batch normalisation, GeLU activation layer and a 1D convolution of kernel size 1 and filter size 125. A skip connection is added between the first 1D convolutional layer and the output of the first layer to feed the subsequent attention layer of filter size 125. After this first block, the following sequential layers are repeated 5 times: batch normalisation, GeLU activation layer, 1D convolution layer with skip connection and kernel size 7, padding 3 and filter size 125, batch normalisation, GeLU activation, 1D convolution kernel size 1 and filter size 125. The output is concatenated with the skip connection and feeded to an attention pooling layer. Finally, there is a global max pooling layer over the position dimension, followed by a dense layer that reduces the input size of 125 to one, passed through a GeLU activation layer. The model was implemented in PyTorch (v2.1.1).

### Model training and evaluation

We divided the 30,607 promoters into six equally sized folds, ensuring that overlapping promoters were assigned to the same fold. One fold was reserved for final model assessment, while the remaining five folds were rotated in a cross-validation fashion. This process involved using four folds for training and one fold for determining the best hyperparameters.

We employed a Poisson negative log-likelihood loss function to account for the non-uniform distribution of the promoter activity score, following a similar approach as Basenji2^59^ and Enformer^8^. The neural network was trained for a total of 10 epochs with a batch size of 128. To optimise the training process, we utilised the AdamW optimizer^60^, with a gradual warm-up learning rate starting from 2 × 10^−8^ which increased linearly for the first 5,000 training batches until reaching 10^−4^. Subsequently, the learning rate was decreased using a cosine function until the last epoch. Other hyperparameters involved were: weight decay = 0.2, *β*_1_= 10^−5^, and *β*_2_= 10^−5^. The model was trained on one NVIDIA Quadro RTX 6000 GPU.

### Generating active synthetic promoters

To generate synthetic human promoters with PARM, we used a genetic algorithm (GA) previously reported by^17^ with slight differences. Using the DEAP-ER framework (v2.0.0, https://github.com/aabmets/deap-er), we set the population size to 200, where each individual was a randomly generated sequence of 232bp. We set the mutation probability to 0.8, the two-point crossover probability to 0.5, and let it run for 500 generations. We used the standard tournament selection operator (sel_tournament) with three individuals participating in each round (contestants=3), and the evaluation operator to be the PARM’s prediction for the K562 model. After running five independent instances of the GA, we cloned 20 synthetic sequences into barcoded plasmids as described above.

### In silico mutagenesis validation library

The activity of each fragment was calculated by taking the mean of the three barcodes to which each individual fragment was linked. To choose a set of promoters and cell lines that were measured reliably we chose a stringent cutoff based on autocorrelations. For each position, we computed the effect of all mutations as a mean of all mutations. Then, we computed the Pearson correlation of each position with the following one, removing the first and last positions. If that autocorrelation with equal to or higher than 0.5, we kept that promoter and cell line set for the following analysis. This resulted in a set of 12 cell line - promoter combination for which we have reliable data.

### Motif insertion library

The activity of each fragment was calculated by taking the mean of the five barcodes to which each individual fragment was linked. Also, we required each fragment to be linked to at least four barcodes and have a pDNA (input) count of > 1000. These requirements led to a final 6 promoters with YY1 insertions, 17 for NFYA, 18 for NRF1 and 14 for SP1.

### Cell lysate preparation for mass spectrometry analysis

Crude nuclear extracts (NE) were prepared as described before^61^. Briefly, cells were collected and spun down (400g, 5 min, 4 °C) then the obtained cell pellet was washed twice with ice-cold PBS. The cell pellet was resuspended in 5 volumes of buffer A (10 mM HEPES KOH pH 7.9, 15 mM MgCl2, 10 mM KCl) and incubated for 10 min on ice. Cells were pelleted by centrifugation (400g, 5 min, 4 °C) and resuspended in 2 volumes of freshly made buffer A+ (buffer A supplemented with 0.15% NP-40 and EDTA-free complete protease inhibitor). Then, cells were lysed by dounce homogenization (40 strokes, 30s rest on ice every 10 strokes), and crude nuclei were collected by centrifugation (3200g, 15 min, 4 °C). Crude nuclei were washed with ice-cold PBS (10 times the volume of the pellet) and clean crude nuclei were collected by centrifugation (3200g, 5 min, 4 °C). Crude nuclei were resuspended in buffer C (420 mM NaCl, 20 mM HEPES pH 7.9, 20% (v/v) glycerol, 2 mM MgCl2, 0.2 mM EDTA, supplemented with 0.1% NP-40, CPI, and 0.5 mM DTT) and incubated for 90 min while rotating at 4 °C. Afterward, the nuclear lysate was centrifuged (20,000g, 30 min, 4 °C) and the soluble nuclear fraction was collected. Obtained NE were aliquoted, snap-frozen in liquid nitrogen, and stored at −80 °C.

### DNA Pull-Downs

DNA oligonucleotides were ordered from Integrated DNA Technologies (IDT) with 5′-biotinylation of the forward strand. The sequences of the oligonucleotides TMEM promoter 70 bp and TMEM 30 bp promoter used can be found in Table S1. Oligonucleotides were annealed as described previously^62^. DNA pull-downs were performed in duplicate as described previously^33^. For DNA pull-downs using stable isotope labeling, experiments were performed in technical duplicate as described previously and experiments were performed three times using NE collected in different days. Similarly, For label free quantification experiments were done in quadruplicate using NE collected in different days. Briefly, for each reaction, 20 μL of streptavidin− sepharose bead slurry (GE Healthcare) was equilibrated by washing once with 1 mL of PBS containing 1% NP-40 and twice with 1 mL of DNA binding buffer (DBB; 1 M NaCl, 10 mM Tris pH 8.0, 1 mM EDTA, 0.05% NP-40). Next, 500 pmol of previously annealed DNA oligonucleotides were incubated with the beads in 600 μL DBB final volume (30 min of rotating at 4 °C), beads were then washed twice with DBB and once with protein incubation buffer (PIB; 150 mM NaCl, 50 mM Tris pH 8.0, CPI, 0.25% NP-40, 1 mM DTT). Per pull-down, 500 μg of NE was incubated with the beads in a total volume of 600 μL PIB (90 min rotating at 4 °C). After the incubation beads were washed three times with 1 mL of PIB and twice with 1 mL of PBS. Excess PBS was removed using a syringe and beads were immediately resuspended in 50 μL of elution buffer (2 M urea, 100 mM Tris pH 8.5, 10 mM DTT) and incubated for 20 min while shaking at room temperature. Samples were then incubated with iodoacetamide (50 mM final concentration) for 10 mins in the dark while shaking. Next, proteins were digested using 0.25 μg of trypsin and incubated while shaking for 2 h at room temperature. Afterwards, samples were spun down, and the supernatant was collected. Beads were washed once more with 50 μL of elution buffer, and the supernatant was collected and added to the previously collected supernatant, 0.1 μg of trypsin was added to the mix and incubated overnight. The next day, samples were purified using StageTips. Dimethyl labelling on StageTips was done as described previously^63^.

### Mass spectrometry analysis

Peptides were analysed by LC-MS/MS on an Orbitrap Exploris 480 mass spectrometer connected to an Evosep One LC system (Evosep Biotechnology, Odense, Denmark). Before LC separation with the Evosep One, peptides were reconstituted in 0.1% formic acid and 20% of the sample was loaded on Evotip Pure™ (Evosep) tips. Peptides were eluted directly on the column and separated using the pre-programmed “Extended Method” (88 min gradient) on an EV1137 (Evosep) column fused with an EV1086 (Evosep) emitter. Nanospray was achieved using the Easy-Spray NG Ion Source (*Thermo*) with spray voltage set to 1.7 - 2 kV. The Exploris 480 was operated in DDA Cycle Time mode with 1 sec. duty cycle and full MS scans being collected in the Orbitrap analyzer with 60,000 resolution at m/z 200 over a 375-1500 m/z range. The default charge state was set to 2+, the AGC target was set to “standard” for both MS1 and ddMS2 and for both scan types the maximum injection time mode was set to “auto”. Monoisotopic peak determination was set to “peptide” and dynamic exclusion was set to 20 sec. For ddMS2, a normalised HCD collision energy of 30% was applied to precursors with a 2+ to 6+ charge state meeting a 5 × 10^4^ intensity threshold filter criterion. Precursors were isolated in the Quadrupole analyzer with a 1.2 m/z isolation window and MS2 spectra were acquired at 15,000 resolution in the Orbitrap.

### Mass spectrometry data analysis

All raw mass spectrometry spectra were processed using Maxquant software v2.4.9.0 (Max Planck Institute of Biochemistry) and searched against the UniProt curated human proteome database released in 2022. Identified proteins were filtered for common contaminants. Dimethyl-labelled samples were analysed using a modified workflow that is based on the built-in dimethylation 3plex method. For quantification, light-dimethyl-labelled peptides (+28.031 Da) and heavy-dimethyl-labelled peptides (+32.056 Da) were used. Low abundance resampling was enabled. Only proteins that were quantified in all 4 channels were used for downstream analysis. Outlier statistics were used to identify significant proteins. Proteins were considered significant with one interquartile range for both forward and reverse experiments. For label free quantification experiments protein identification and relative abundance quantification was done using Maxquant software v2.4.9.0 using Andromeda as search engine. Statistical analysis was done using Perseus v2.0.1.1. Results were filtered for contaminants, reverse hits and only proteins that were found in all samples were considered. Volcano plot strategy was used to identify potential interactors combining t-test P value with ratio information.

### In silico saturated mutagenesis (ISM)

As described by many others ^6,8,19,23^, ISM involves predicting the promoter activity of a reference sequence and sequences with all possible single mutations at the position of interest. The importance score is calculated as the score of the reference sequence minus the average score of all possible mutations at a single position.

### ICGC data

International Cancer Genome Consortium (ICGC) data was obtained from IGCG data portal release 28^29^, querying whole genome sequencing (WGS) as the sequencing strategy and retrieving all donors with data in chromosome 5, from 1,295,152 to 1,295,462 bp in the hg19 reference genome.

### Enformer predictions

For the Enformer comparisons, we predicted the variant effect as previously described by Karollus et al.^54^. We computed the CAGE outputs for the relevant cell types and log-transformed the predictions. We averaged both strands, the three neighboring bins around the SNP positions, and all available CAGE tracks for the relevant cell lines. For missing cell-types such as LnCaP, we used the best matching alternatives PC3 and DU145.

### Prediction of the direction of effect of cis-eQTLs

To evaluate the accuracy of the predicted expression effect of the PARM model we compared the predicted direction of effect with the sign of the measured beta of fine-mapped GTEx v8 cis-eQTLs^30^. The concordance in the direction of effect is defined as:

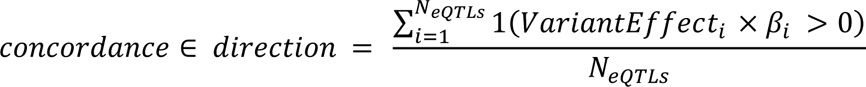

Where the *N*_*eQTLs*_ indicates the total number of eQTLs (1836) and *β* indicates the observed eQTL effect as estimated in GTEx. As a baseline comparison, we computed the concordance in direction for random predictions by randomly sampling from a uniform distribution using the *random.uniform* implementation from NumPy set from −1 to +1^64^.

### Motif scanning and RS identification

Using the Hocomoco v11 human database^21^, we scanned the importance scores of the 30,607 promoters generated by one of the (cell type-specific) PARM models. Each of these 30,607 vectors, containing promoter importance scores, were scanned with each Information Content (IC) motif from the Hocomoco database with a stride of 1. At each position, we computed the Pearson correlation coefficient (PCC) between the IC of the motif and the importance score at each position, here referred to as *r*_*position*_. Moreover, we calculate the mean of the effect scores multiplied by the IC, here referred to as *IS*_*position*_.

We only retain hits with a *r*_*position*_ equal to or higher than 0.75. To avoid hits in regions where the importance score was low but still had a PCC higher than 0.75 (which occurred especially for short and high-content motifs), we applied a second cutoff on *IS*_*position*_ that accounts for a motifs’ varying length and information content, as follows:

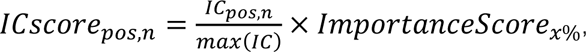

where *IC*_*pos*,*n*_ indicates the information content of a motif in a specific position *pos* and nucleotide *n*, and *max*(*IC*) is the maximum value of the *IC* across all positions and nucleotides over that motif. The *ImportanceScore*_*x*%_ represents the xth percentile value of all variant effect scores in all promoters (described above). Thus, *ICscore*_*pos*,*n*_ would represent the minimal motif to consider a hit, where the most important position would have a value of *ImportanceScore*_*x*%_. Then, the cutoff of *IS*_*position*_ is defined as:

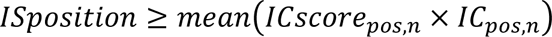

The *ImportanceScore*_*x*%_ is calculated separately for each model due to differences in the range of importance scores between models, as well as separately between activators and repressors. For activators, the 95th percentile is used, while for repressors we consider the bottom 5th percentile.

Thus, we consider an RS if a motif in a database survives both cutoffs in *r*_*position*_ and *IS*_*position*_ in the in silico mutagenesis profile.

### Identifying unknown motifs

To identify RS that did not match any known motif, we scanned for patches of at least 5 nucleotides where all their importance scores were equal to or higher than the *ImportanceScore*_95%_ for activators or equal to or lower than the *ImportanceScore*_5%_ for repressors. If there were several consecutive RS separated by a maximum of two nucleotides, they were treated as a single RS. Then, the nucleotide importance of the full RS region was correlated with all motifs in the Hocomoco v.11 human^21^ or the Jaspar2024 CORE vertebrates non-redundant database^32^. If neither database contained a hit with PCC≥ 0.75, it was annotated as an ‘unknown motif’.

### Clustering motifs to obtain motif families

To cluster motifs separately for (1) the Hocomoco v11 human database, and for (2) the unknown motifs, we followed the methods outlined in Vierstra et al^65^. Pairwise similarity scores of Position frequency matrices (PFM) were computed using TOMTOM (v5.4.1) with a minimum overlap of 6 bp and Pearson correlation as the distance metric for scoring alignments. Subsequently, we converted these scores to E-values by a log transformation, after adding a pseudocount of 1 × 10^−8^. Hierarchical clustering was then conducted using complete linkage and Pearson correlation as the distance metric. The tree was cut at a height of 0.8, and all motifs within the cutoff were considered from the same motif family.

### Matching RS with its TF expression

For further analysis, we exclusively included RS associated with expressed Transcription Factors (TFs). Expression status was determined using data from the Human Protein Atlas version 23.0, accessible at https://www.proteinatlas.org/download/rna_celline.tsv.zip^31^. RS linked to TFs for which holds that:

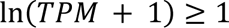

were selected for subsequent analysis.

### Consensus motif

We selected the motif with the lowest E-value as the representative motif to construct the consensus motif within a cluster in the unknown motifs. For each motif in the cluster, its Position Frequency Matrix (PFM) was added to the representative motif, considering the optimal offset and orientation reported by TOMTOM. Each position was then normalised by the number of motifs added in that position. If a position in the consensus motif was contributed by less than 30% of the motifs in the cluster, that position was removed.

### Sequence scanning

We performed a naive scan of all promoter sequences spanning a region from 250 bp upstream to 50 bp downstream of the TSS using FIMO (v5.4.1)^20^ with default arguments and the Hocomoco v11 human database^21^.

### RS insertion analysis

We performed in silico insertion of YY1, SP1, NRF1, and NFYA motifs in all promoters. For each of these TFs, we inserted three variants: the consensus sequence, and two mutated versions as negative controls. We shortened the consensus sequence as defined by Hocomoco v11^21^ to work in similar motif length range. For NRF1, consensus sequence: “ACTGCGCATGCGCA”, mutated sequence 1: “ACTGCGTATTCGCA”, mutated sequence 2: “ACTTCGAATTGGCA”. For NFYA, consensus sequence: “TCAGCCAATCAGAA”, mutated sequence 1: “TCAGTCTATCAGAA”, mutated sequence 2: “TCATCAGATCAGAA”. For SP1, consensus: “GGGGGCGGGGCCGG”, Mutated 1: “GGGGCCCGGGCCGG”, Mutated 2: “GGAGGTGGTGCCGG”. For YY1, consensus: “CAAGATGGCGGC”, mutated sequence 1: “CAAGTTGCCGGC”, mutated sequence 2: “CACGAAGACGGC”. The motifs were inserted in 15 positions ranging from 160 bp upstream of the TSS to 63 bp downstream in a fixed step of 14 nt.

### Endogenous promoter activity

To get an estimate of endogenous promoter activity in K562, we aggregated the following datasets:

- GRO-cap from^66^: GEO Series accession number GSM1480321 (https://www.ncbi.nlm.nih.gov/geo/query/acc.cgi?acc=GSM1480321)
- csRNA from^67^: GEO Series accession number GSE135498 (https://www.ncbi.nlm.nih.gov/geo/query/acc.cgi?acc=GSE135498)
- NetCage from^68^: GEO Series accession number GSM3318243 (https://www.ncbi.nlm.nih.gov/geo/query/acc.cgi?acc=GSM3318243)
- Stripe-Seq from^69^: GEO Series accession number GSM4231304 (https://www.ncbi.nlm.nih.gov/geo/query/acc.cgi?acc=GSM4231304)

We obtained the raw files (fastq files) and utilised the FASTX Toolkit 0.0.14 with default parameters to trim 20 nucleotides from the end of each read. Subsequently, we employed Bowtie2 2.5.1 with default settings to align the reads against the human rDNA (U13369.1). Reads that did not align to the rDNA were then mapped to the hg19 reference genome. We generated bigwig files using bamCoverage 3.5.2, setting a minimum mapping quality of 30, an offset of one, and a bin size of one. These bigwig files were then overlaid with our set of 30,607 TSS. Finally, we conducted Principal Component (PC) analysis and used the first component (PC1) as a combined measure of endogenous promoter activity, which explains 94.19% of the variance.

---

**Supplementary Table1.**
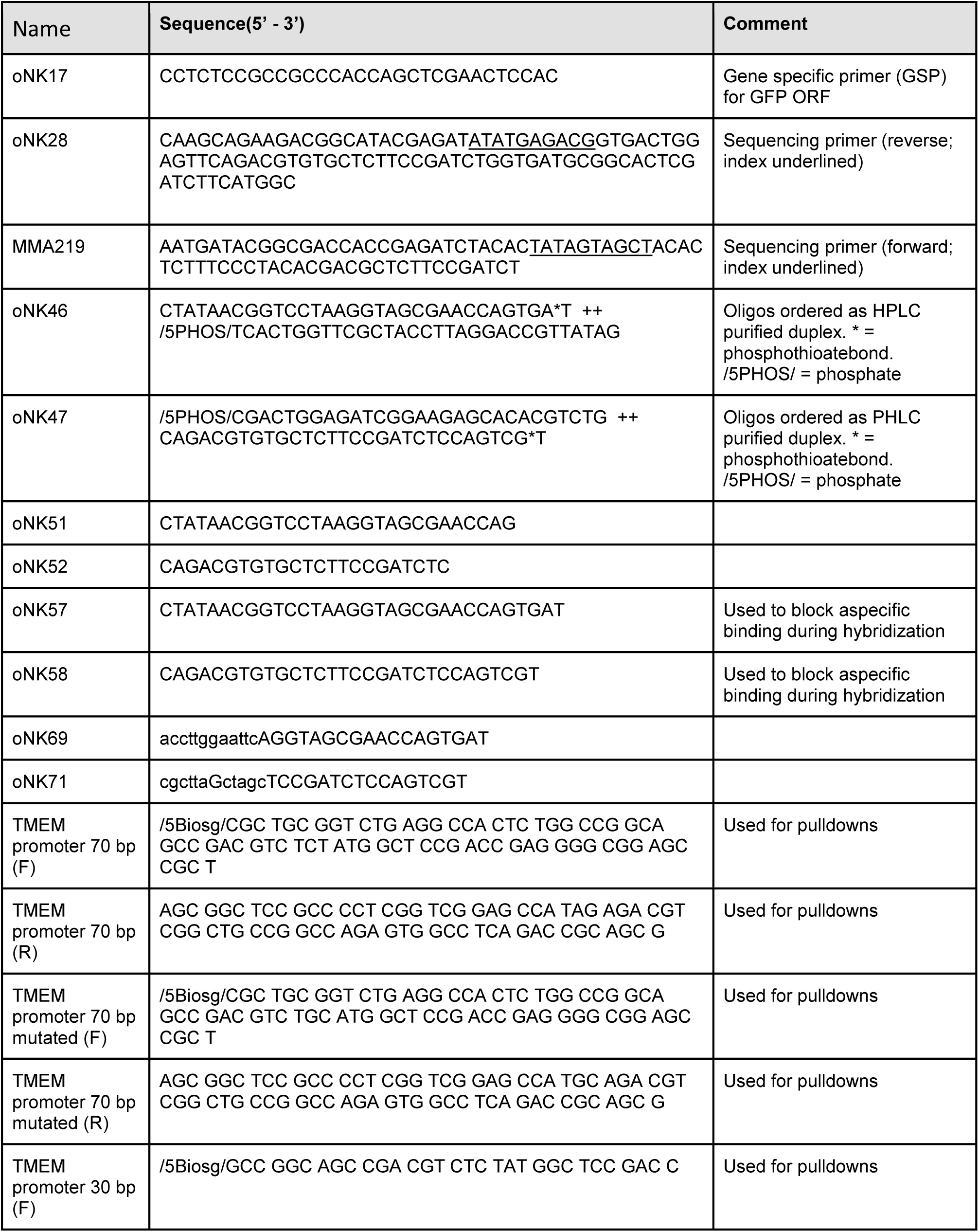
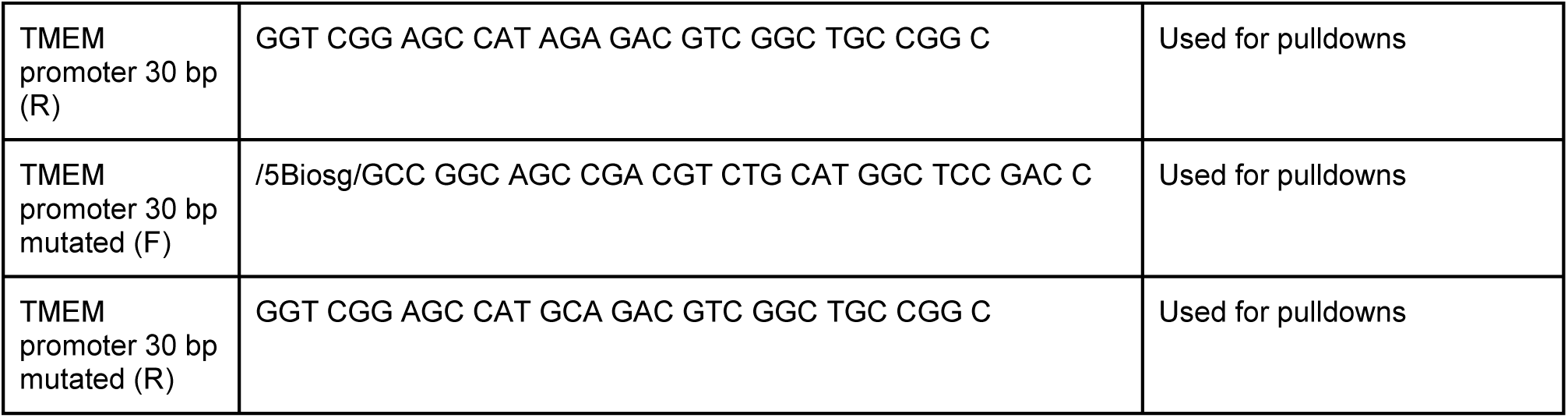
Oligonucleotide sequences.

## DATA AND CODE AVAILABILITY

All raw sequencing data generated will be made publicly accessible upon acceptance of the manuscript. The code used for prediction and training PARM can be found in https://github.com/vansteensellab/PARM. The in-silico mutagenesis plots of all 30,607 promoters for the 5 cell types as described in **Figure 5** can be found in https://surfdrive.surf.nl/files/index.php/s/jmMqlxX8LrnqgHM and will be made publicly available.

## ACKNOWLEDGMENTS

This work is part of the PERICODE project. We thank Maryam Akbarzadeh, Sarah Derks, Michiel Thiecke and members of our laboratories for stimulating discussions; the AvL Foundation and NKI Genomics, Proteomics, and Research High-Performance Computing core facilities for support; Max Trauernicht for providing MPRA vector. Co-funded by the European Union (ERC, RE_LOCATE, 101054449 to B.v.S.). Views and opinions expressed are however those of the author(s) only and do not necessarily reflect those of the European Union or the European Research Council. Neither the European Union nor the granting authority can be held responsible for them. Research at the Netherlands Cancer Institute (NKI) is supported by an institutional grant of the Dutch Cancer Society and of the Dutch Ministry of Health, Welfare and Sport. The Oncode Institute is partially funded by the Dutch Cancer Society.

## SUPPLEMENTARY FIGURES

**Figure S1:**
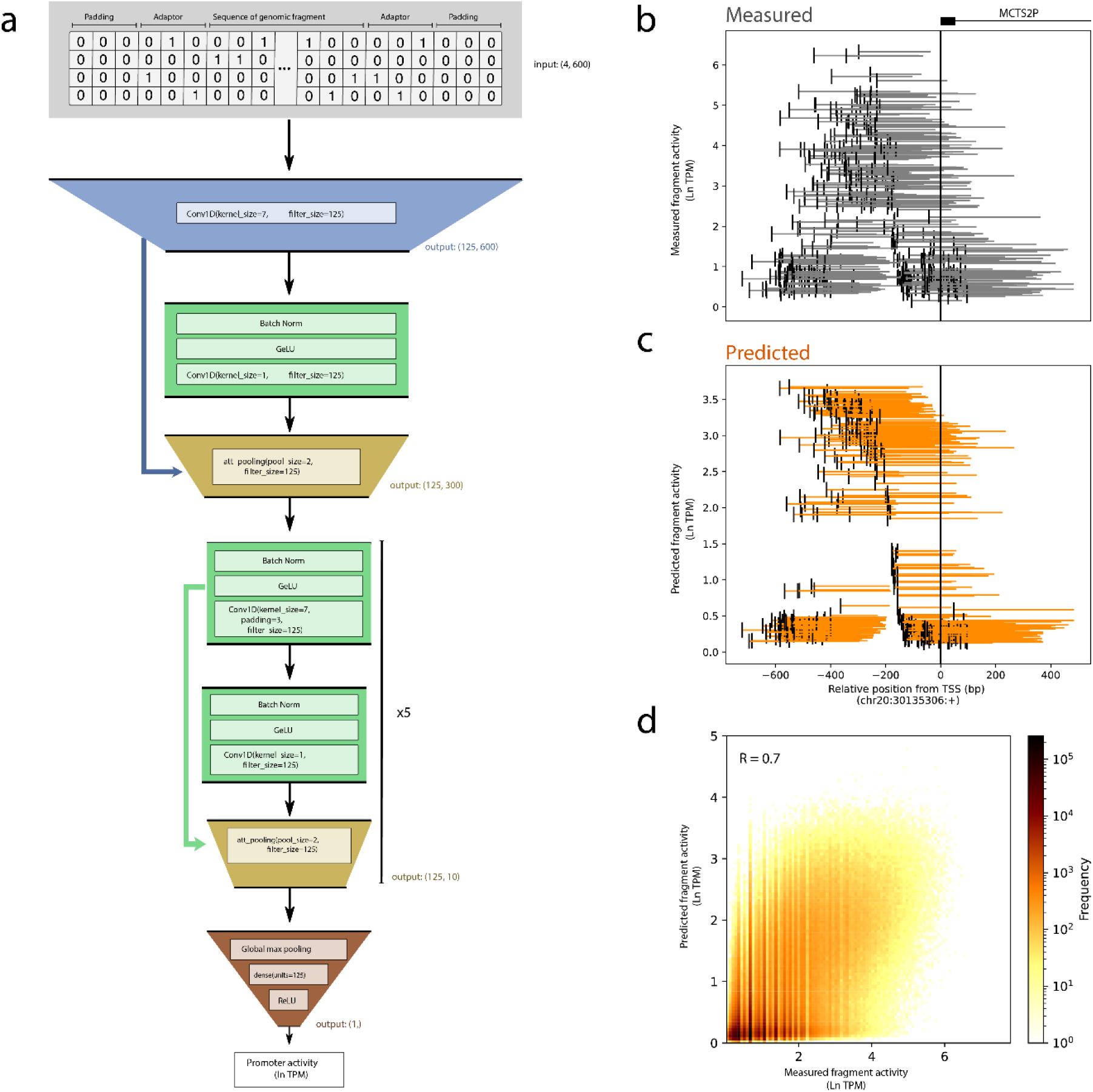
PARM architecture. **a,** Deep learning architecture of PARM (see Methods), right integers indicate the output shape. PARM takes as input a one-hot encoded sequence of the MPRA fragment. Adaptors are added to the sequence and differ between genome-wide library and focused library. **b-c,** Fragments overlapping the MCTS2P promoter in the sense orientation and their corresponding measured (**b**) and predicted (**c**) fragment activity. Note that measurements are noisier when compared to predictions. The black bar indicates the 5′ end of each fragment. **d,** Correlation between measurements and predictions at fragment level in the test set for K562.

**Figure S2.**
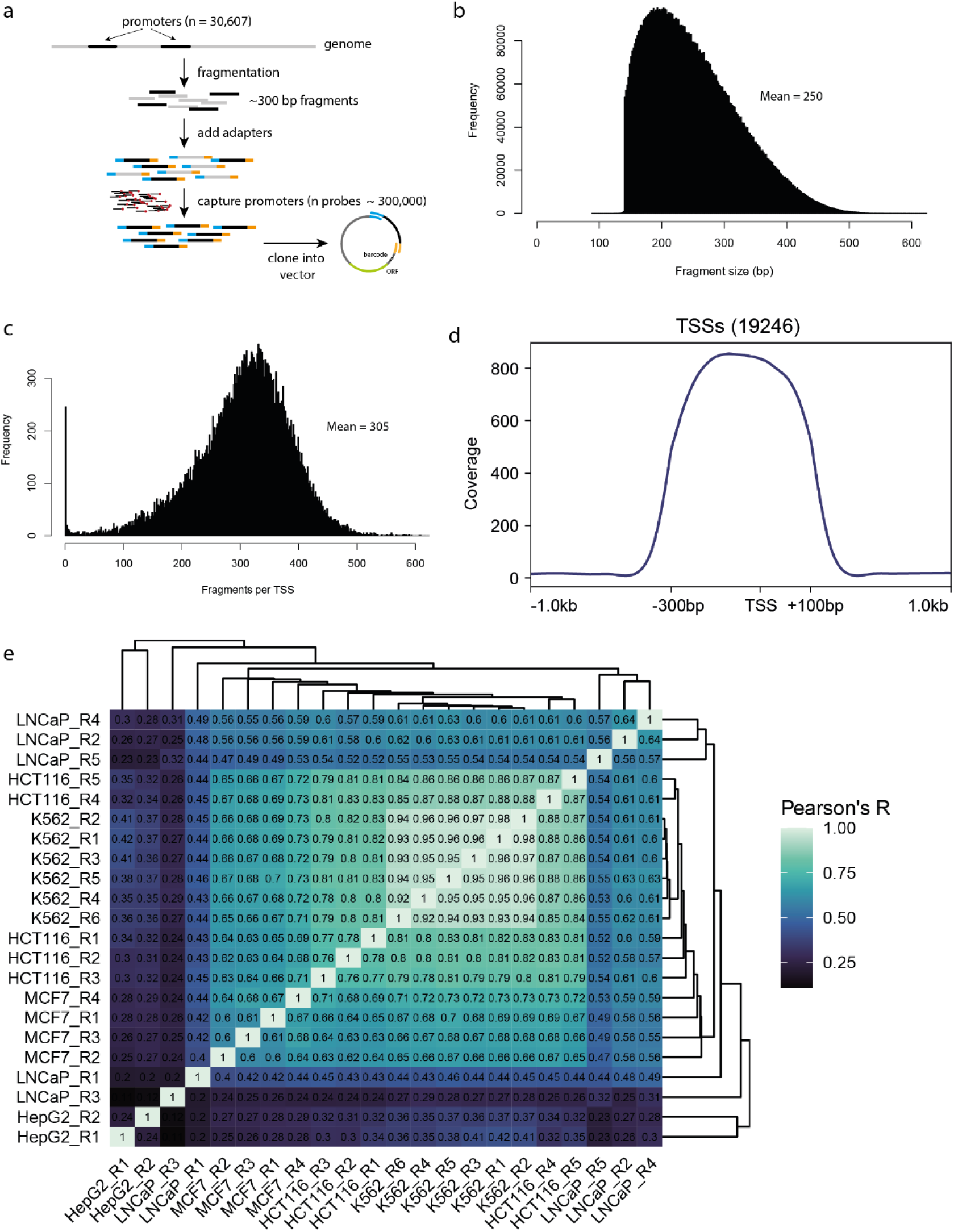
Focused-library design and characterization. **a,** Outline of the construction of a promoter-focused MPRA library. See Methods for details. **b,** Distribution of fragment sizes in the promoter-focused library. **c,** Number of fragments per TSS in the focused library. **d,** Average fragment coverage per position relative to the TSS, shown for 19,246 TSS that are at least 3 kb away from any other TSS. **e,** Correlations between measured promoter focused library data (averaged by TSS) of independent replicate experiments.

**Figure S3.**
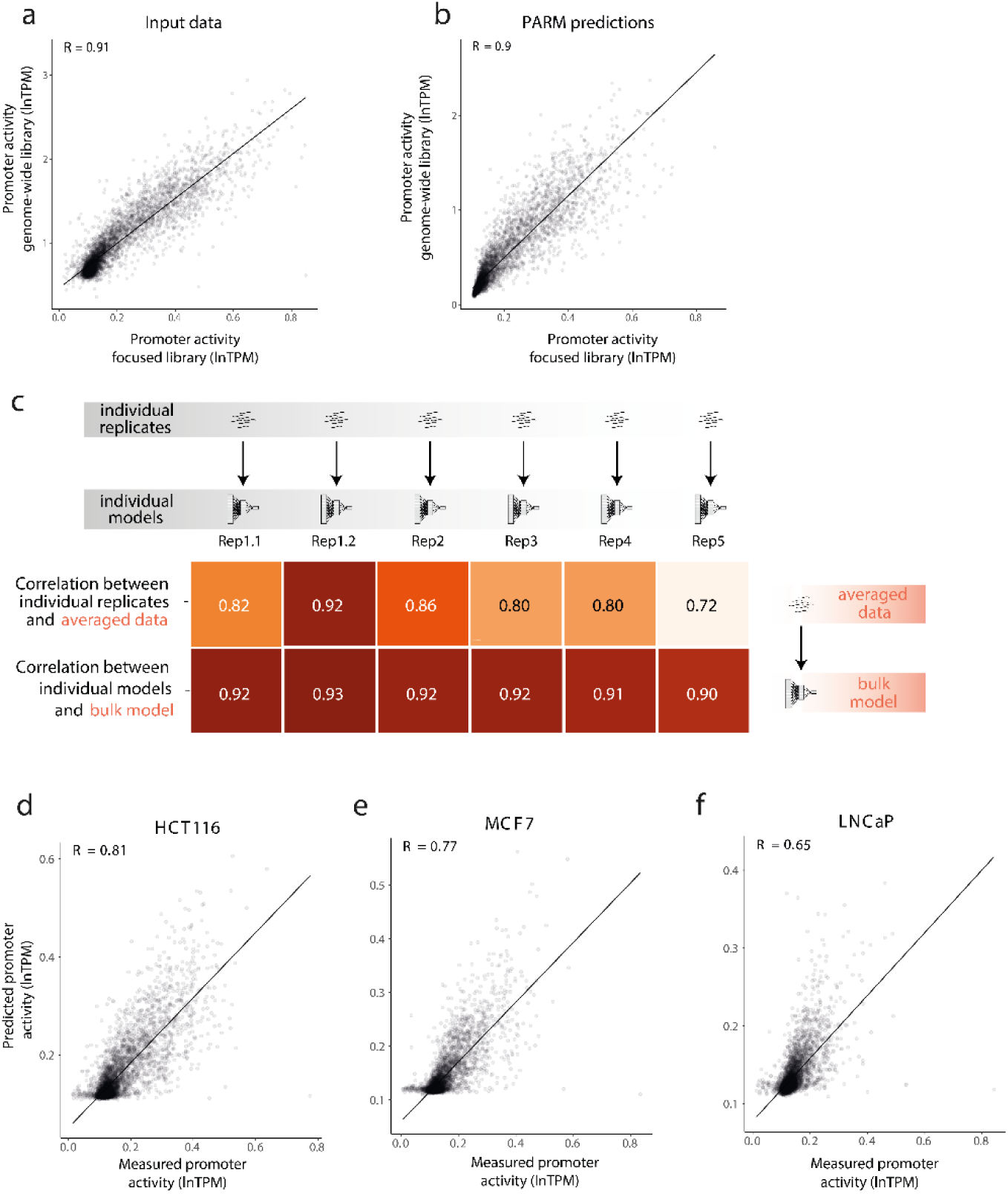
Promoter-focused library MPRA to train PARM. **a,** Strong correlation between promoter activities in K562 cells measured by MPRA using either full-genome libraries (data from ∼1.6 billion cells^18^) or the promoter-focused library (data from ∼50 million cells; this study).. **b**, Strong correlation between PARM-estimated promoter activities after training on either full-genome and focused MPRA data as in **a**.. **c,** Potent noise suppression by PARM. Models trained on individual replicate experiments correlate better and more consistently with a model trained on all replicates combined (bottom data row), than the corresponding individual replicates of input MPRA-data correlate with their average (top data row). **d-f,** Correlations between PARM-predicted and measured activity of promoters based on MPRA data obtained with promoter-focused libraries. Plots in **a, b, c, d-f** show correlations for 17% of promoters that were left out from the training data.

**Figure S4.**
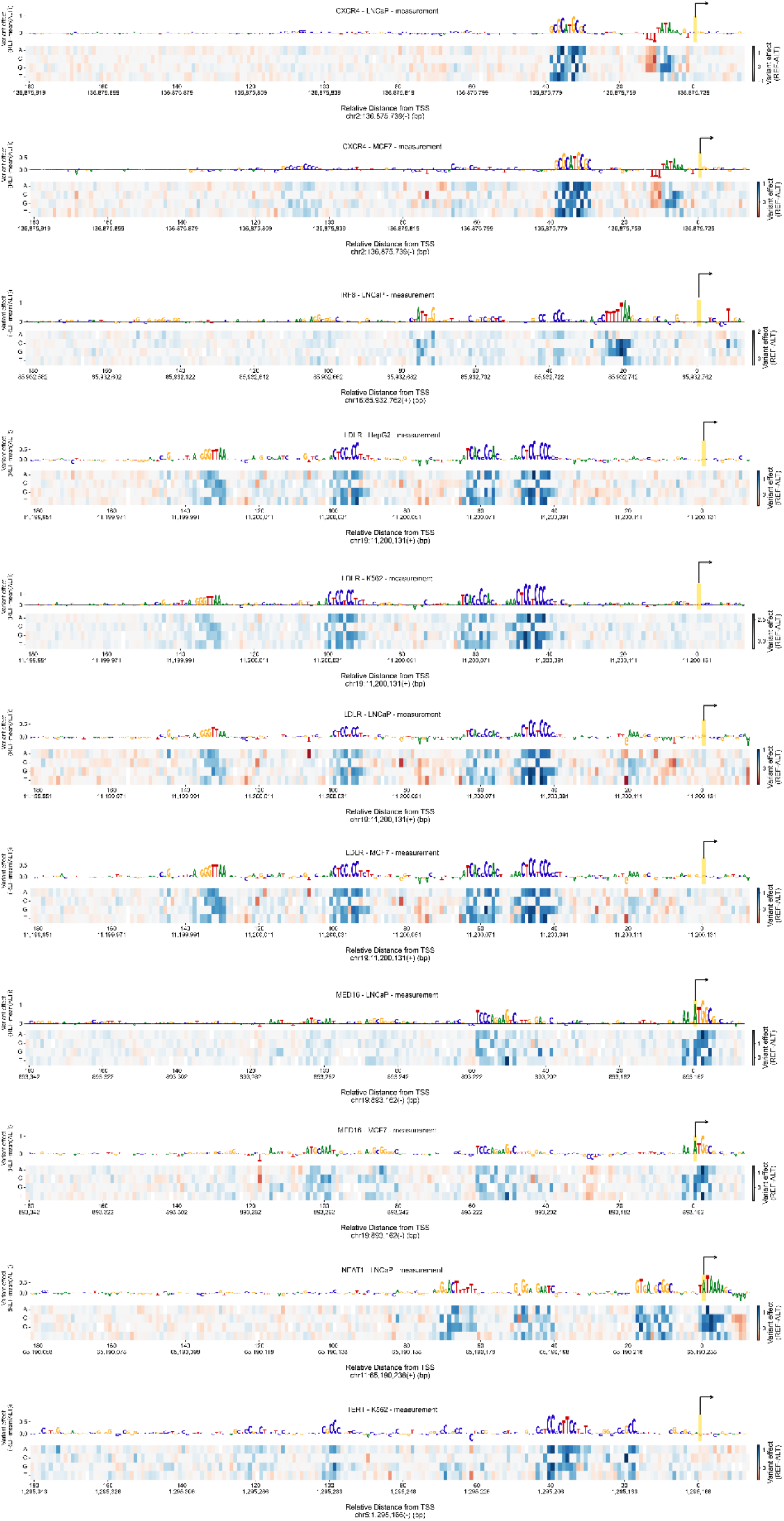
*In vitro* mutagenesis plots of all promoters and cell line combinations displayed in Figure 3d.

**Figure S5.**
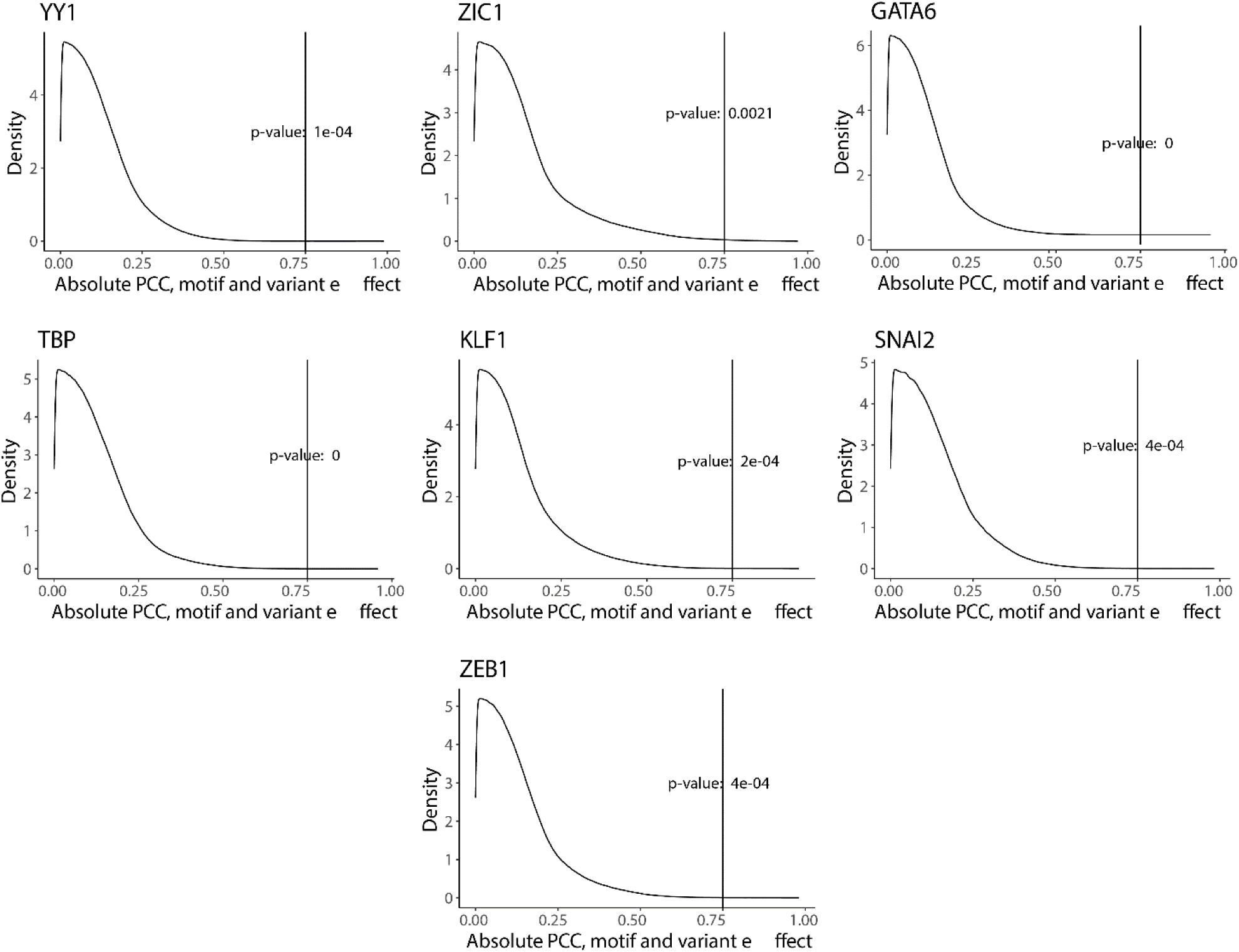
Distribution of Pearson correlation coefficients (PCC) of indicated TF motifs was assessed by scanning over 30,607 randomly shuffled promoters. Each promoter was shuffled randomly while maintaining dinucleotide frequencies, and the variant effect was computed using PARM in K562. For each position in the 30,607 shuffled promoters, we computed the PCC between the motifs from Hocomoco v11^21^ and the variant effect. The absolute PCC values are plotted. Notably, the |R| > 0.75 cutoff is rarely achieved in these randomised data.

**Figure S6.**
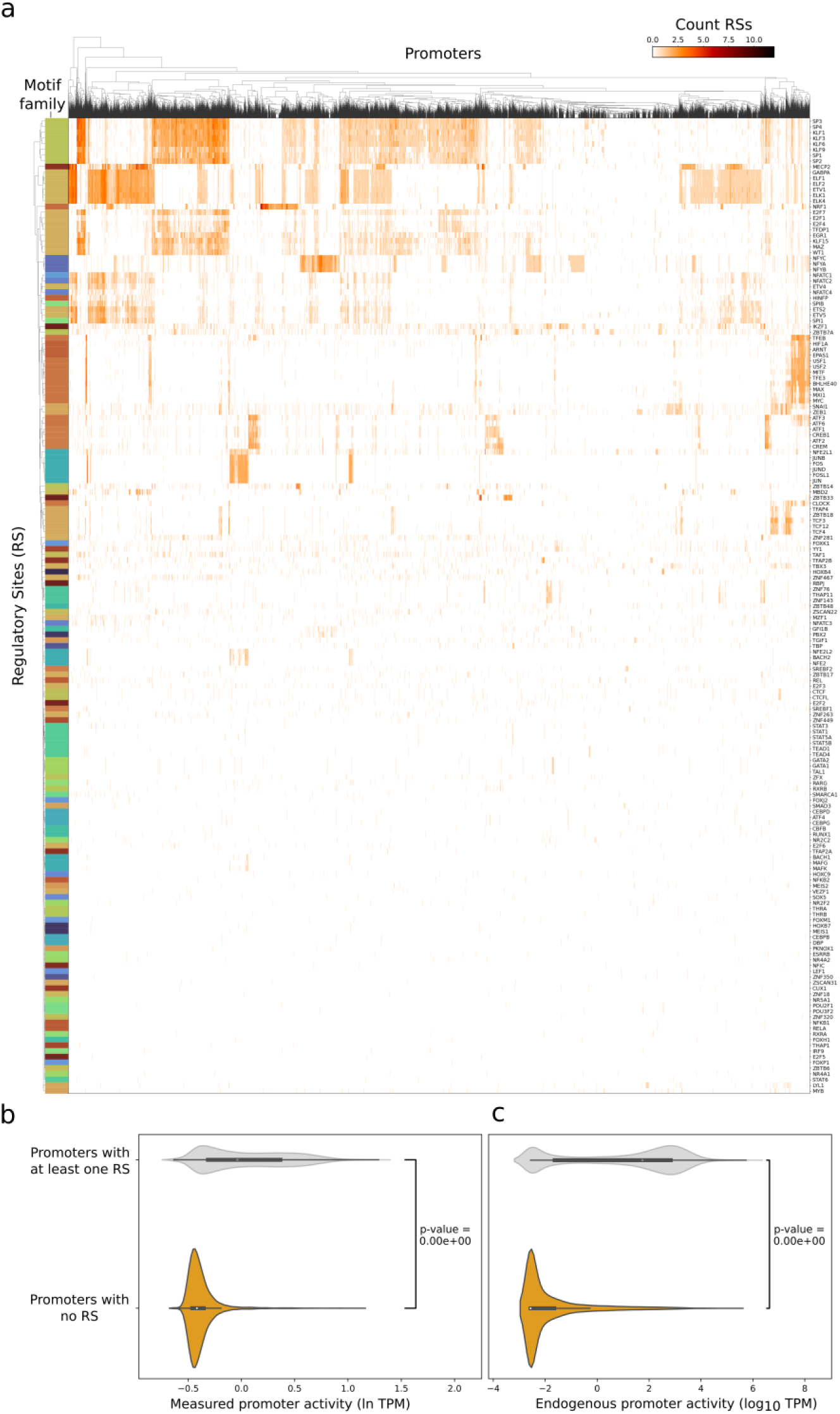
TF motif annotation of all RSs in K562 cells; correspondence with promoter activities. **a,** Full matrix of TF motifs that match RSs across 20,543 promoters with at least one RS. Colours in the column on the right depict motif families (see Methods). Hierarchical clustering was performed using complete linkage and euclidean distance as metric. **b,** Promoters with at least one RS are generally more active than those without any RS, according to (left panel) MPRA data^18^ and according to (right panel) a combination of measures of endogenous promoter activity (PC1, see Methods).

**Figure S7.**
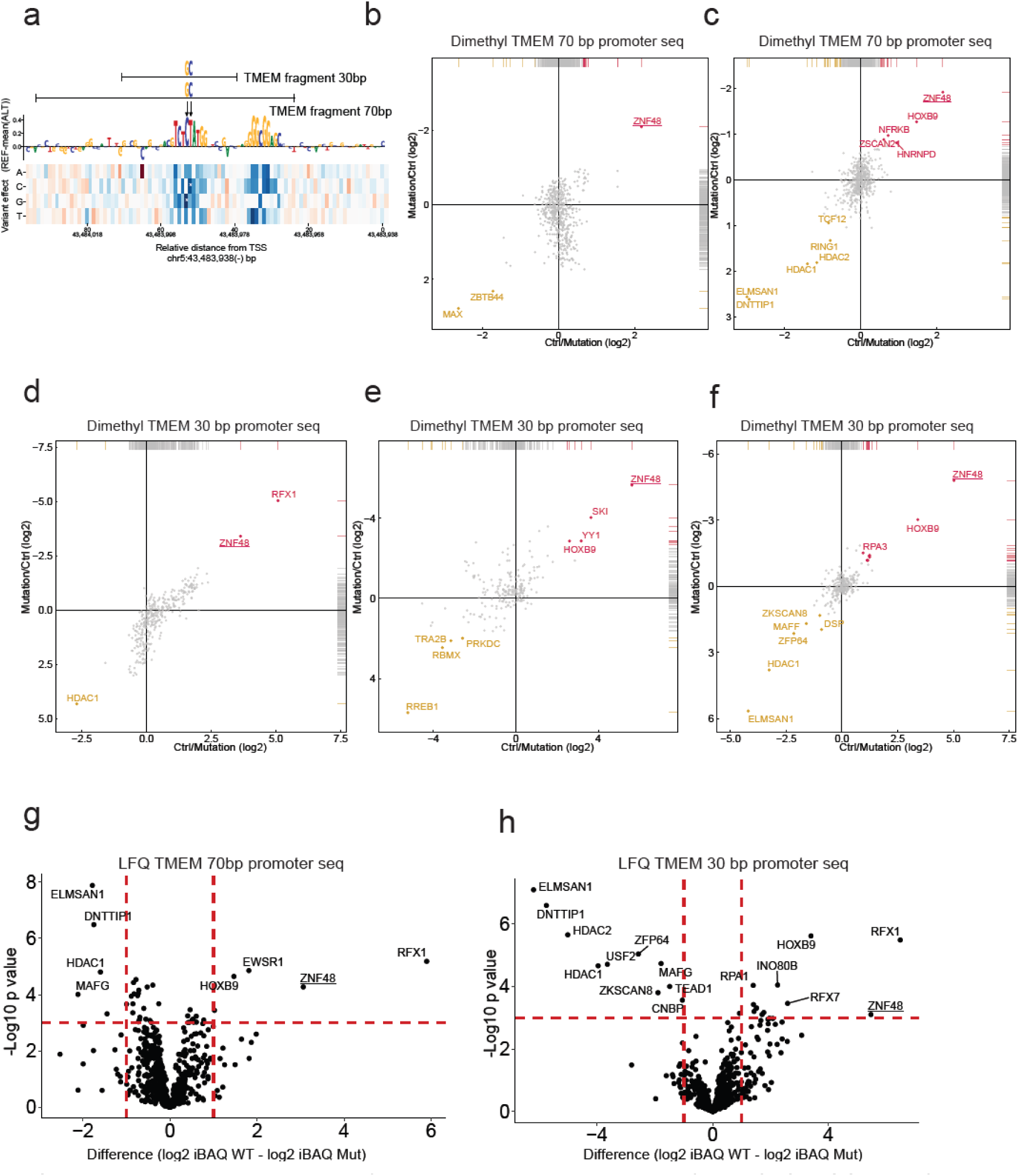
Interaction proteomics identifies ZNF48 as potential binder of the TCTCATGGT motif. **a,** schematic representation of the oligo baits used for isotope-based and label free-based DNA pull downs of TMEM 70 bp promoter and TMEM 30bp promoter sequence (for full oligo sequence see Table S1). **b-c,** biological replicates DNA pulldown followed by dimethyl labeling of the TMEM 70 bp promoter sequence. Each experiment was done in duplicate with label swap of replicates. Enriched proteins that bind preferentially to the TMEM 70 bp promoter sequence (WT control) are labeled magenta and proteins labeled yellow showed preferential binding to the mutant sequence (Mut). **d-f,** three biological replicates of DNA pulldown of the TMEM 30 bp promoter sequence followed by dimethyl labeling. **g-h,** Label free quantification mass spectrometry analysis of proteins captured by DNA pulldown of the TMEM 70 bp promoter (**g**) or TMEM 30 bp promoter sequence (**h**) compared to mutant sequence. Label free quantification of quadruplicates was used and proteins were considered significant with a p-value <0.001 and a log2 intensity based absolute quantification (iBAQ) value fold-change of 1.

**Figure S8.**
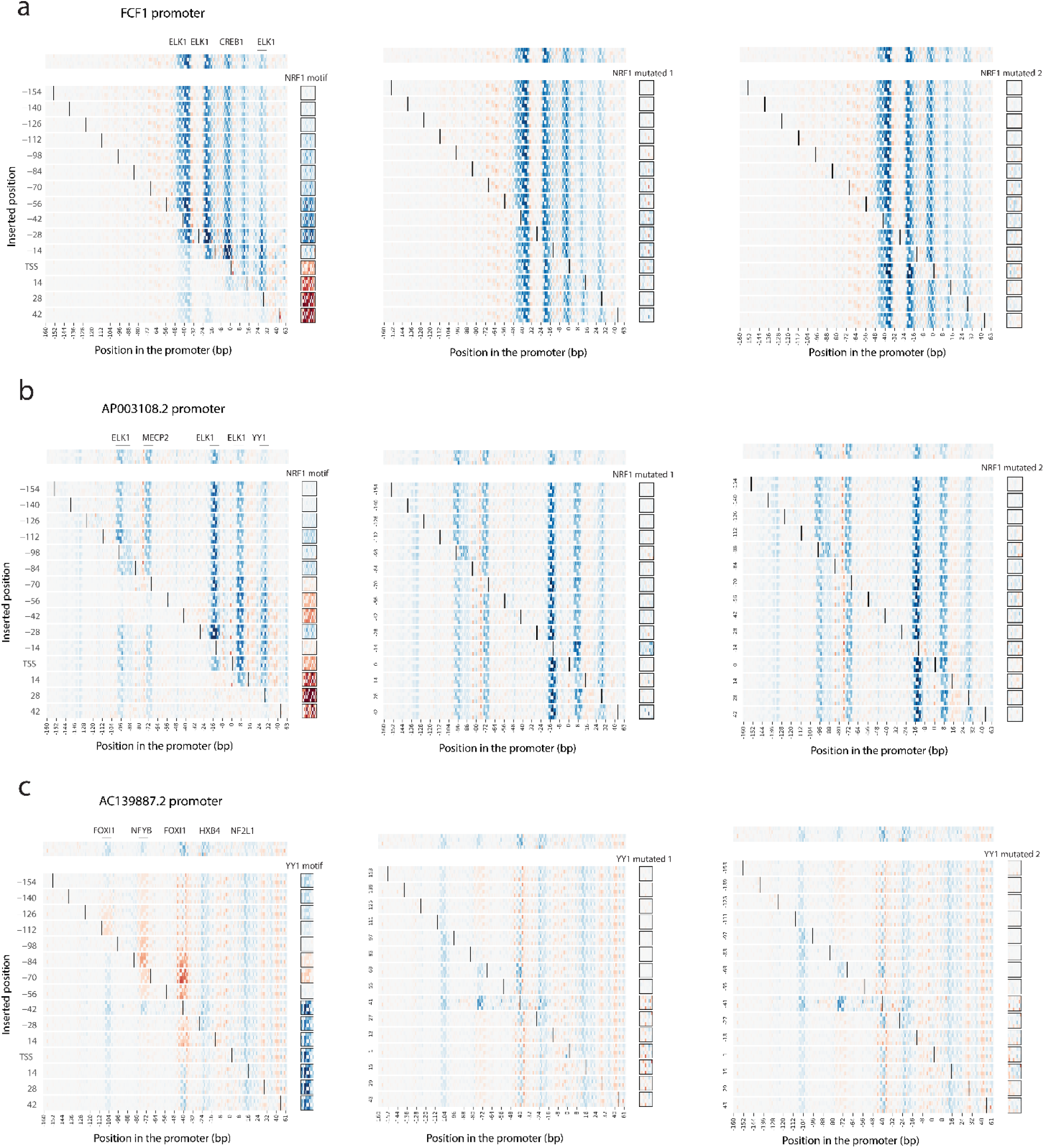
Predicted effects of TF motif insertions are specific. **a-c**, Left column shows PARM-predicted effects of insertion of NRF1 (a, b) and YY1 (c) motif insertions. Middle and right columns show the same analyses with mutated motifs that are predicted to abolish TF binding (see Methods). Note that the mutated motifs are virtually inactive.

